# Integrating genetic dependencies and genomic alterations across pathways and cancer types

**DOI:** 10.1101/2020.07.13.184697

**Authors:** Tae Yoon Park, Mark D.M. Leiserson, Gunnar W. Klau, Benjamin J. Raphael

## Abstract

Recent genome-wide CRISPR-Cas9 loss-of-function screens have identified genetic dependencies across many cancer cell lines. Associations between these dependencies and genomic alterations in the same cell lines reveal phenomena such as oncogene addiction and synthetic lethality. However, comprehensive characterization of such associations is complicated by complex interactions between genes across genetically heterogeneous cancer types. We introduce SuperDendrix, an algorithm to identify differential dependencies across cell lines and to find associations between differential dependencies and combinations of genetic alterations and cell-type-specific markers. Application of SuperDendrix to CRISPR-Cas9 loss-of-function screens from 554 cancer cell lines reveals a landscape of associations between differential dependencies and genomic alterations across multiple cancer pathways in different combinations of cancer types. We find that these associations respect the position and type of interactions within pathways with increased dependencies on downstream activators of pathways, such as *NFE2L2* and decreased dependencies on upstream activators of pathways, such as *CDK6*. SuperDendrix also reveals dozens of dependencies on lineage-specific transcription factors, identifies cancer-type-specific correlations between dependencies, and enables annotation of individual mutated residues.

## Introduction

A key problem in cancer biology is to identify the genes that cancer cells depend on for their growth and survival advantage. This knowledge both informs our understanding of cancer development and also suggests therapeutic targets [1, 2, 3]. While some of these cancer-essential genes are altered by somatic mutations and thus identified by high-throughput DNA sequencing [4, 5, 6], other genes are rarely or not mutated in cancer, such as lineage-specific transcription factors or master regulators [7, 8, 9, 10]. An alternative approach to identify cancer dependencies is to perturb genes in *in vitro* cancer models, such as cell lines, and measure growth or viability following such perturbations. Genes whose perturbation results in a change in viability reveal potential cancer-specific gene dependencies. Recent technologies such as genome-wide pooled RNAi [11] or CRISPR [12, 13] loss-of-function screens enable high-throughput genome-wide perturbation screens. Projects such as DRIVE [14] and the Cancer Dependency Map (DepMap) [15, 16] have applied these technologies to hundreds of cancer cell lines and identified genes that are essential in specific cancer cell lines, often referred to as conditionally essential genes, or differential dependencies. Combining differential dependencies with genomic mutations identified in the same cell lines has revealed several context-specific dependencies including examples of oncogene addiction [17, 18] and synthetic lethality [19, 20].

Recently, several studies have attempted to systematically identify associations between differential dependencies and genomic alterations using data from large-scale RNAi and CRISPR datasets [14, 16, 21, 22, 23, 24, 25, 26]. The computational approaches used in these studies can be grouped into two classes. The first class of approaches attempt to identify associations between differential dependencies and *single* genomic biomarkers [14, 16, 21, 22, 23, 24]. These approaches recapitulate many of the classic oncogene addiction and synthetic lethal interactions, as well as a few additional associations. However, the restriction to a single biomarker limits the statistical power to detect associations with rare genomic alterations occurring in a small subset of cell lines. It is well known that driver mutations in cancer exhibit a “long tail” with relatively few frequently mutated genes and many rarely mutated genes [4, 5] of driver mutations in cancer. Thus, it is reasonable to assume that rare genomic alterations may also be associated with genetic dependencies. One explanation for the long tail of driver mutations is that many of the functions perturbed in cancer involve interactions between many genes in pathways and networks [5, 27, 28]. A genetic dependency may similarly be associated with genomic alterations in many genes, and thus examining single alterations or single genes may miss many important associations.

The second class of computational approaches attempt to identify associations between differential dependencies and sets of *multiple* biomarkers [15, 25, 26]. For example, Tsherniak et al. [15] used a random forest classifier to predict dependencies in the DepMap dataset from genomic alterations. While thousands of associations are reported, many of the reported associations are with gene expression markers and other frequent events, which is not surprising since the classifier skews toward explaining the most frequent associations. REVEALER [25] leverages the observation that driver mutations in the same pathway tend to be mutually exclusive across tumors, i.e., that few tumors have more than one driver mutation in a given pathway [29, 30, 31]. This method generalizes earlier approaches that identify sets of mutually exclusive mutations in cancer genomes [31, 32, 33, 34, 35], by focusing the search for sets of mutually exclusive mutations to cell lines with the the highest dependency scores. However, REVEALER’s iterative search does not scale to the analysis of thousands of gene perturbations and genomic alterations [26]. Furthermore, REVEALER does not evaluate statistical significance of the reported associations, leading to many false positive associations. Recently, UNCOVER addressed the scalability issues of REVEALER using a combinatorial optimization approach [26]. Unfortunately, when applied to large-scale RNAi or CRISPR datasets, UNCOVER predicts hundreds-to-thousands of associations, and the quality of such predictions is generally unknown. These issues are due to three factors. Spurious associations are a significant challenge in analyzing large-scale RNAi or CRISPR screens, since the number of phenotypes (gene perturbations) and number of features (genomic alterations) are orders of magnitude larger than the number of samples. This challenge is further exacerbated when searching for sets of multiple biomarkers as the number of such sets is massive and the optimal set is unknown *a priori*.

We introduce a new algorithm, SuperDendrix, to identify sets of approximately mutually exclusive genomic alteration and cell type features that are associated with differential dependencies from large-scale perturbation experiments. SuperDendrix includes several key features: (1) a principled approach to identify and score differential dependencies using a 2-component mixture model; (2) a combinatorial model and optimization algorithm to find feature sets associated with differential dependencies; (3) a model selection criterion to select the size of the associated set and a robust statistical test that accounts for different frequencies of genomic alterations across samples.

We apply SuperDendrix to identify associations between genotypes and differential dependencies in a large-scale CRISPR-Cas9 screen dataset from DepMap of 17,634 gene knockouts across 558 cancer cell lines from 31 cancer types. We identify 32 associations between sets of alterations and differential dependencies, and an additional 117 associations that include cancer-type features. Many of these associations group into well-known cancer pathways including the NFE2L2, RB1, MAPK, and Wnt pathways. We observe that associations between differential dependencies and genomic alterations within a pathway respect the topology and regulatory logic of the interactions within the pathway. Specifically, we find that cell lines containing oncogenic mutations in a gene upstream in a pathway (either activating mutations in an oncogene or inactivating mutations in a tumor suppressor genes) often have *increased dependencies* on genes downstream in same pathway. These associations generalize the phenomenon of oncogene addiction to *oncogene pathway addiction* [17, 18]. On the other hand, we find that oncogenic mutations in genes that are downstream in a pathway are often associated with *decreased dependency* on genes upstream in the same pathway. SuperDendrix also identifies associations between differential dependencies and cancer types, most prominently dependencies involving lineage-specific transcription factors and a previously unreported association between *TCF3* dependency in multiple myleoma or B-cell lymphomas with mutations in *ID3* or *MEF2B*.

SuperDendrix is a rigorous computational approach to identify associations between gene dependencies, genomic alterations, and cell-type features in large-scale RNAi and CRISPR screens. SuperDendrix’s high specificity focuses experimental efforts on a small number of the most promising associations. SuperDendrix software is publicly available at https://github.com/raphael-group/superdendrix and results on DepMap datasets are available through a web portal https://superdendrix-explorer.lrgr.io/.

## Results

### SuperDendrix

We introduce SuperDendrix, an algorithm to identify sets of binary features such as genomic alterations and/or cell types that are (approximately) mutually exclusive and associated with a quantitative phenotype. SuperDendrix is generally applicable to any quantitative phenotype, but in this work we focus on the phenotype of cell viability change following genome-wide CRISPR-Cas9 loss-of-function screens. The inputs to SuperDendrix are the following.

- Cell viability measurements from a genome-wide CRISPR-Cas9 loss-of-function screens. We represent these measurements in a phenotype matrix *P* where each entry *p_gj_* of *P* the indicates viability of cell line *j* when gene *g* is knocked out. Each of these scores quantify the *dependency* of a cell line on a gene. We refer to the dependency scores for a gene across cell lines (i.e. row of *P*) as a *dependency profile*.
- A list of somatic alterations in each cell line. Here, we analyze somatic missense and nonsense mutations.
- (Optionally) Categorical information (e.g. cell type) of each cell line.

SuperDendrix consists of three modules (Fig. 1a): (1) Scoring differential dependencies and selecting genomic and cell-type features. (2) Finding feature sets associated with differential dependencies. (3) Evaluating the statistical significance of associations. We briefly describe these three modules below. Further details are in Methods.

**Figure 1:**
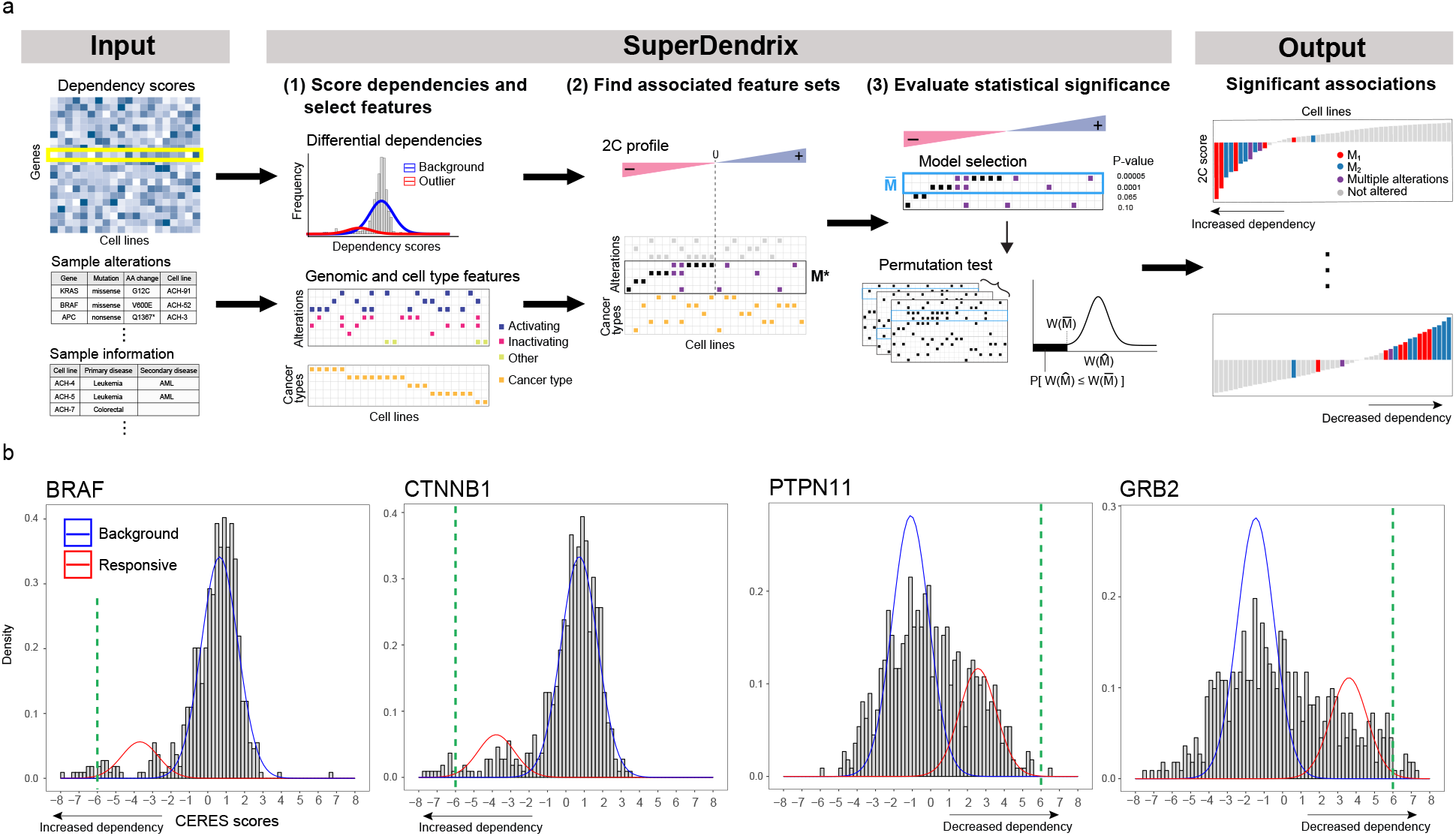
Overview of SuperDendrix. (a) SuperDendrix inputs are dependency scores of gene knockouts from CRISPR-Cas9 screens, genomic alterations, and optionally, cell-type features. In the first step, SuperDendrix scores *differential dependencies* – genes whose dependency scores are better fit by a mixture distribution of two components – and also constructs a genome alteration and cell-type feature matrix. In the second step, SuperDendrix finds a subset *M\** of features that maximize the SuperDendrix weight *W* (*M*). In the third step, SuperDendrix performs model selection to define a subset 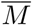 of features that substantially contribute to weight, and computes statistical significance of weight 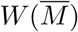 using a permutation test. Associations with false discovery rate (FDR) ≤ 0.2 are output, and include associations between features and increased dependency on profile (top right) and between features and decreased dependency on features (bottom right). (b) Examples of 2C differential dependencies from DepMap data that result from fitting the dependency scores with a mixture model. Blue curve is the background component, and red curve is the responsive component. Green dashed lines indicate 6*σ* criterion of Tsherniak et al. [15], which identifies only a subset of cell lines that are responsive to knockout. *BRAF* and *CTNNB1* are examples of profiles with increased dependency in response to knockout while *PTPN11* and *GRB2* are examples of decreased dependency.

#### (1) Scoring differential dependencies and selecting genomic and cell-type features

The first module in SuperDendrix has two steps: (i) scoring differential dependencies from the dependency scores; (ii) selecting the genomic alteration and cell-type features that will be evaluated for association. In the first step, we derive a *differential dependency profile* for each gene knockout (row of *P*). This profile quantifies the magnitude of the effect on the gene knockout on each cell line relative to a background distribution. We assume that the dependency scores *p_g_*_1_*, …, p_gn_* for knockout *g* are generated from two populations: a *background* population that is unaffected by the knockout, and a *responsive* population that is affected by the knockout. We fit a two-component mixture model to the dependency scores *p_g_*_1_*, …, p_gn_*, and decide whether the score distribution is better fit by one-component or two-components using the Bayesian Information Criterion (BIC). In the case where the two-component fit is preferred, we say that the cell lines are *differentially dependent* with respect to the gene knockout *g*, or that gene *g* is a *differential dependency*. We define the *differential dependency score*, or *2C score*, 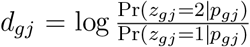 for cell line *j* as the log ratio of the posterior probabilities that cell line *j* is from component 2 (*z_gj_* = 2) and that cell line *j* is from component 1 (*z_gj_* = 1). We choose component 1 to be the component with smaller mean so that negative 2C scores indicate decreased viability, or *increased dependency* in response to knockout. Conversely, positive 2C scores indicate *decreased dependency* in response to knockout. We assume that a minority of cell lines are responsive to gene knockout and thus refer to the component that contains fewer cell lines as the *responsive* component and the component with more cells lines as the *background* component. Summarizing this analysis, we say that differential dependencies whose responsive component has negative scores are *increased dependencies*, and those whose responsive component has positive scores are *decreased dependencies*.

Next, we construct the genomic alteration and cell-type feature matrix *A*. This matrix contains two parts. The first part of *A* consists of genomic alterations. We use the OncoKB database [36] to select genes and mutations with annotated roles in cancer. For each cancer gene in OncoKB, we use the functional annotations of missense and nonsense somatic mutations to create three gene alteration features: missense activating mutations are combined into a single feature labeled `GENE(A)`; missense and nonsense inactivating mutations are combined into a single feature labeled `GENE(I)`; and the remaining unannotated missense mutations are combined into a single feature labeled `GENE(O)`. The second part of *A* contains cell-type features. In this analysis, we construct a binary feature for each cancer type represented in the analyzed cell lines. Each cancer type feature has the value 1 for cell lines of the corresponding cancer type and the value 0 for cell lines of other cancer types. We note that by definition, the cancer-type features are mutually exclusive across cell lines. Further details of both steps are in Methods.

#### (2) Finding feature sets associated with differential dependencies

The second module in SuperDendrix is a rigorous and practically efficient combinatorial optimization algorithm to find sets *M* of features (genomic alterations and/or cell-type features) in *A* that are: (i) approximately mutually exclusive; and (ii) associated with increased (or decreased) dependency. We derive the SuperDendrix weight *W* (*M*) of a set *M* that combines criteria (i) and (ii), and use an integer linear program (ILP) to find the set *M\** of minimum (or maximum) weight *W* (*M\**) (Methods).

#### (3) Evaluating statistical significance of associations

The third module of SuperDendrix includes two steps. First, a *model selection* step identifies a subset 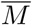 of the features in *M\** found in the second module, where each feature in 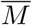 contributes significantly to the weight 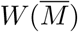. This step uses a conditional permutation test to iteratively remove features whose contribution to the weight *W* (*M\**) is nearly the same as random features. Second, we assess the statistical significance of the set 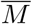 using a permutation test that conditions on *both* the number of cell lines that have each alterations and the number of alterations per cell line. Since the number of genomic alterations varies considerably across cell lines (Fig. S1), controlling for the number of alterations per cell line is important, as previously noted for predicting cancer driver mutations [37, 38].

We also developed an interactive tool for visualization and exploration of the SuperDendrix results which is available at: https://superdendrix-explorer.lrgr.io/. Further details of SuperDendrix as well as comparisons to another recent method [26] for mutual exclusivity analysis are in Methods.

### Identification of differential dependencies and alteration features in DepMap

We used SuperDendrix to analyze the Avana dataset [19Q1/2.19.2019] from Project DepMap containing results of CRISPR-Cas9 loss-of-function screens of 17,634 genes across 558 cancer cell lines from 31 cancer types [15, 39]. DepMap provides a dependency score for each gene knockout across all cell lines. These dependency scores are computed using the CERES algorithm [39], which is designed to reduce confounding at loci with high genomic copy number. CERES scores are scaled across all gene knockouts so that the median score for known “essential” genes is 1 and the median score for genes with “no dependency” is 0. Following the “6*σ*” criterion of Tsherniak et al. [15], 1,730 genes have at least one cell line with a CERES score at least six standard deviations below or above the mean. We refer to CERES score profiles for these 1,730 genes as 6*σ differential dependencies* (Table S1).

The first step of SuperDendrix computes that 492 (28%) of the 6*σ* differential dependencies are better fit by the two-component mixture model. We refer to these genes as *two-component (2C) differential dependencies* (Fig. 1b, Fig. S2, Table S2). The 492 2C differential dependencies include 413 genes with increased dependency and 79 genes with decreased dependency. We find that 92 of the 2C differential dependencies are in the COSMIC Cancer Gene Census (CGC) [40] (*P <* 0.001) – including *BRAF*, *KRAS*, *NRAS*, and *PIK3CA* (Fig. S3) – a higher precision than for non-2C genes (2C: 18.7%, non-2C: 11.0%). These 2C genes are enriched (FDR *<* 0.05) for 122 GO molecular functions [41, 42] including kinase activity, G protein-coupled receptor activity, and DNA binding (Table S3). We find that the 6*σ* differential dependencies that are not 2C differential dependencies either contain only a few outlier samples (e.g. 86% have fewer than 4 outlier samples) or have dependency score distribution that are unimodal with large variance (Fig. S4).

For the second input to SuperDendrix, we use 420,541 non-synonymous single-nucleotide mutations reported in sequencing data from the Cancer Cell Line Encyclopedia (CCLE) [43] for 554 of the 558 cell lines analyzed by DepMap. We also include the annotated cancer type of each cell line as a feature. For the feature selection step in the first module of SuperDendrix, we use mutation annotations from OncoKB [36] and obtain an alteration matrix of 566 alteration features in 363 genes with a total of 9,464 alterations across the 554 cell lines. Further details are in Methods.

### Associations between genomic alterations and differential dependencies

We used SuperDendrix to identify associations between sets of genomic alterations and the 492 2C differential dependencies. SuperDendrix identifies 23 single alterations and 9 sets of approximately mutually exclusive alterations that are significantly associated (FDR ≤ 0.2) with differential dependencies (Fig. 2a and Table S4). 25 of these sets are associated with increased dependency and 7 with decreased dependency. Many of these associations are well-known dependencies including examples of oncogene addiction (e.g. `BRAF(A)` and increased dependency on *BRAF* [47]) and synthetic lethality (e.g. `ARID1A(I)` and increased dependency on `ARID1B` [48]). Half of the associations (16/32) group into three well-known cancer pathways (NFE2L2, RB1, and MAPK), and we highlight novel findings of SuperDendrix in these pathways.

**Figure 2:**
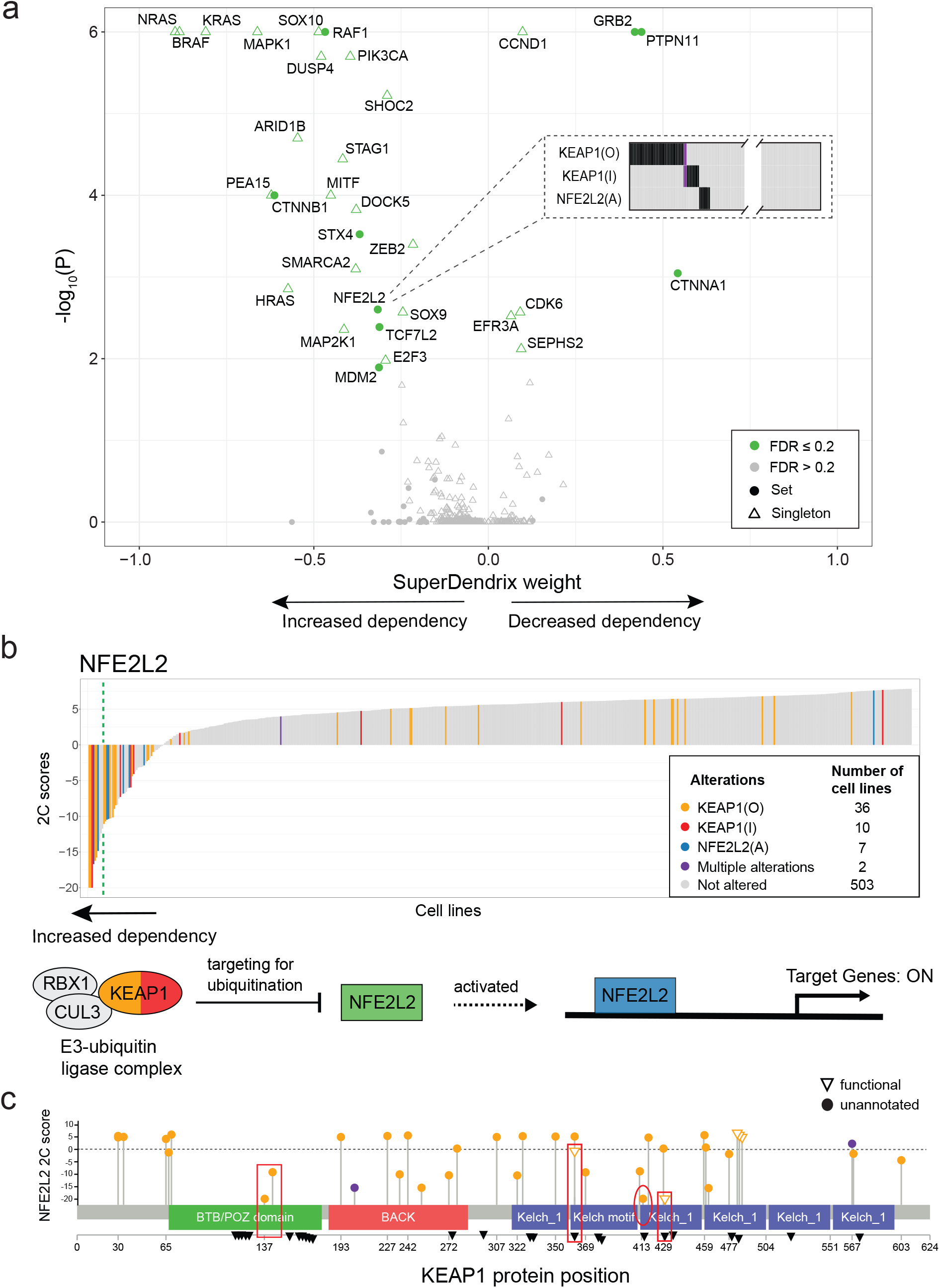
SuperDendrix identifies associations between genomic alterations and 2C differential dependencies in multiple biological pathways. (a) SuperDendrix weights and *P*-values for 26 2C differential dependencies with significant (*FDR* ≤ 0.2) associations with genomic alterations. 8 of these associations are sets of multiple alterations; e.g. the set {`NFE2L2(A), KEAP1(I), KEAP1(O)`} are alterations are (approximately) mutually exclusive (inset) and associated with *increased* dependency on *NFE2L2*. (b) (Top) Waterfall plot of 2C differential dependency scores for *NFE2L2* across cell lines. Cell lines are colored by status in associated alteration set {`NFE2L2(A), KEAP1(I), KEAP1(O)`}. Green dashed line indicates 6*σ* threshold. (Bottom) KEAP1-NFE2L2 pathway. Solid circles are genes on the pathway, with colors indicating their alterations. Green boxes are genes that are knocked out. Association between *KEAP1* inactivating mutations and *increased* dependency on *NFE2L2* is consistent with the role of KEAP1 as an upstream activator of NFE2L2. (c) Locations of missense mutations in KEAP1 protein that are annotated as **O**ther. `KEAP1(O)` mutations associated with increased dependency on NFE2L2 include: two mutations in the BTB/POZ domain (boxed) which is important for dimerization of KEAP1 [44], two annotated mutations in two of Kelch domains (boxed) which mediate interaction with NFE2L2 [45], and one mutation (circled) that lies at a residue that interfaces with NFE2L2 [38]. Orange (resp. purple) amino acid changes are in cell lines with exclusive (resp. multiple) mutations in *KEAP1*. Triangles indicate locations of mutations that are reported in Uniprot [46] to affect KEAP1-NFE2L2 interaction.

First, SuperDendrix finds an association between the set {`NFE2L2(A)`, `KEAP1(I)`, `KEAP1(O)`} of three alterations and increased dependency on *NFE2L2* (Fig. 2b). The KEAP1-NFE2L2 pathway is frequently perturbed in cancer with activating mutations in *NFE2L2* or inactivating mutations in *KEAP1* reported in more than 30% of lung squamous tumors [49, 50]. `NFE2L2(A), KEAP1(I)`, or `KEAP1(O)` alterations occur in 51/554 of the DepMap cell lines including 31% (4/13) of lung squamous cancer cell lines. Moreover, three alterations are nearly mutually exclusive with only 2/51 altered cell lines having more than one alteration (Fig. S5). Increased dependency on *NFE2L2* in cell lines with `NFE2L2(A)` alterations is consistent with the oncogene addiction model [17, 18], since *NFE2L2* is an oncogene in various cancers including lung, pancreas, breast, and gall bladder [50, 51]. The increased *NFE2L2* dependency in cell lines with *KEAP1* inactivating mutations is consistent with KEAP1’s role in inhibiting NFE2L2 by targeting NFE2L2 for degradation via ubiquitination. Inactivation of KEAP1 results in translocation of NFE2L2 to the nucleus where NFE2L2 targets over 200 genes for transcription [44, 52]. Thus, the increased dependency on *NFE2L2* in cell lines with *KEAP1* inactivating mutations can be viewed as an another form of oncogene addiction, and *NFE2L2* addiction in cell lines with somatic mutations in *KEAP1* or *NFE2L2* was recently reported [53]. Note that only a fraction of the associated alterations occur in cell lines whose CERES score is below the 6*σ* threshold (Fig. 2b), demonstrating the advantages of SuperDendrix’s 2C differential dependency score.

We find that associations with differential dependencies are also useful for annotating individual missense mutations. Several of the `KEAP1(O)` alterations – which include missense mutations that are unannotated in OncoKB – occur in cell lines with strong evidence of increased dependency on *NFE2L2*. These include 2 missense mutations in *KEAP1* (G364C, G430V) that are not reported as functional in OncoKB, but occur at positions that are reported to disrupt NFE2L2 repression [45]. Another two mutations are in the BTB/POZ domain, a domain that is important for KEAP1 dimerization and KEAP1-CUL3 binding [44] (Fig. 2c). Furthermore, the only `KEAP1(O)` mutations in the BTB/POZ domain are two mutations with negative 2C scores: the 21 `KEAP1(O)` with positive 2C scores mutations all lie outside the domain (*P* = 0.07; binomial test). These findings prioritize these mutations for functional validation studies.

Second, SuperDendrix identifies associations between *RB1* inactivating mutations and differential dependencies in *CCND1*, *CDK6*, and *E2F3*, three members of the RB1 pathway (Fig. 3). Cell lines with *RB1* inactivating mutations have *increased dependency* on *E2F3*. It is known that active RB1 binds and inhibits E2F3 transcription factor activity, and that dissociation of the RB1-E2F3 complex results in E2F3-initiated transcription of target genes that promote G1/S transition [54]. Our results suggest that cell lines with inactivating mutations in *RB1* become highly dependent on E2F3 activity, a phenomenon analogous to oncogene addiction [17, 18]. On the other hand, we observe that cell lines with *RB1* inactivating mutations are associated with *decreased dependency* on *CCND1* and on *CDK6*. The CCND1-CDK4/6 complex is known to inactivate RB1 by phosphorylation. Thus, it makes sense that cell lines with inactive *RB1* do not require CCND1 or CDK6 to inactivate RB1, making these cell lines *less* sensitive to knockout of CCND1 and CDK6. This result suggests a correspondence between the direction of dependencies and the boolean logic in a pathway: there is increased dependency on an inhibitor of transcription factor *E2F3*, but decreased dependency on inhibitors of an inhibitor of the transcription factor *E2F3*.

**Figure 3:**
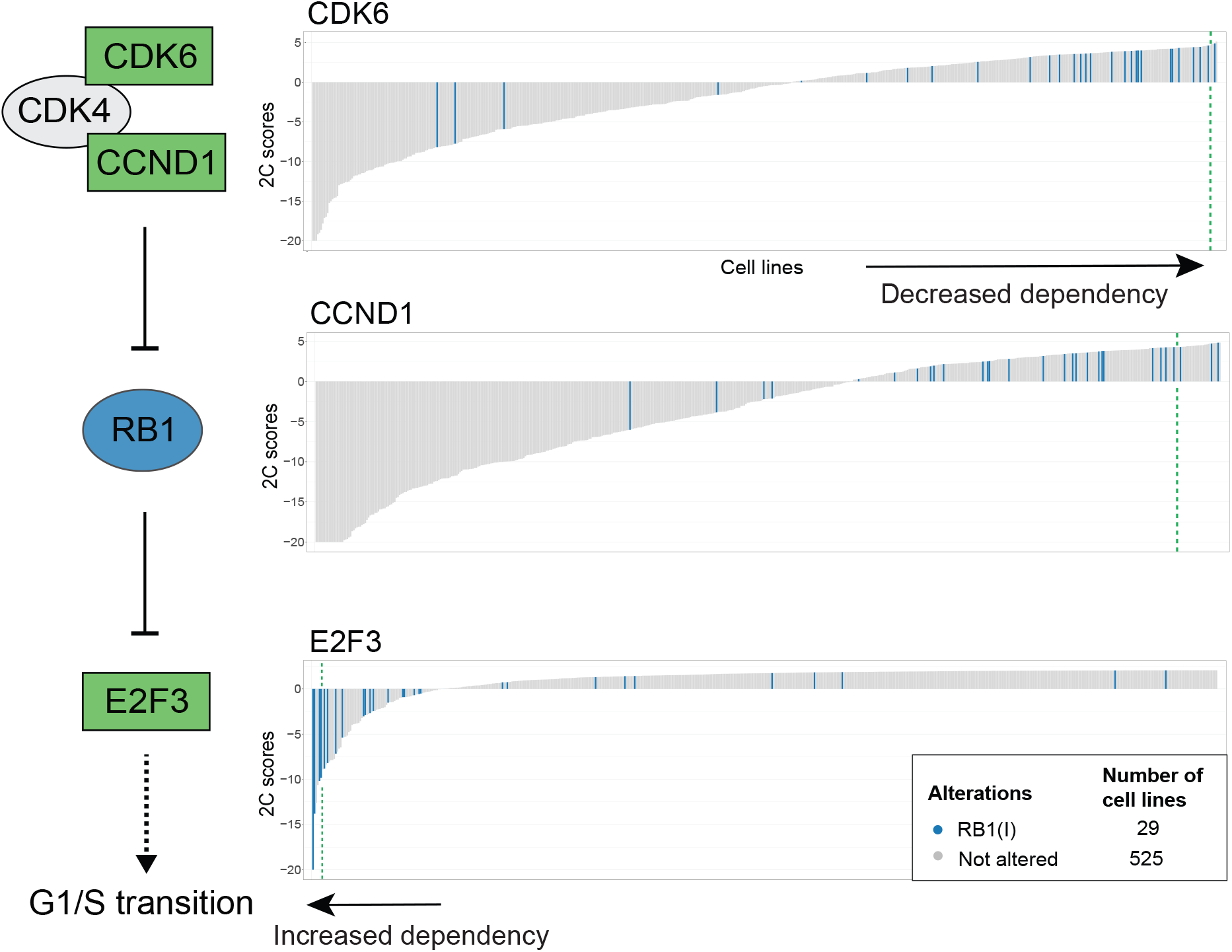
SuperDendrix identifies associations between genomic alterations and 2C differential dependencies in the RB1 pathway. *RB1* inactivating mutations are associated with *increased* dependency on *E2F3*, consistent with RB1’s role in inactivating the E2F3 transcription factor (Same format as Fig. 2b). On the other hand, *RB1* inactivating mutations are associated with *decreased* dependency on *CDK6* and *CCND1*. This is consistent with the role of the CDK4/6-CCND1 complex in inactivating RB1.

Third, Superdendrix finds associations between 12 differential dependencies in the MAPK pathway and subsets of the approximately mutually exclusive alteration set {`BRAF(A), KRAS(A), NRAS(A), HRAS(A), MYD88(O)`} (Fig. 4a). These include well-known associations between activating mutations in *BRAF*, *KRAS*, *NRAS*, or *HRAS* and increased dependency on the corresponding gene [47, 55, 56, 57]. We also find an association between increased dependency on *RAF1* and the set {`KRAS(A), NRAS(A)`} of approximately mutually exclusive alterations. This is consistent with the role of RAF1 as a mediator of RAS for signal transduction in the MAPK pathway during transformation [58]. We also find an association between increased dependency on *SHOC2* and `NRAS(A)` alterations as reported previously [59]. SuperDendrix also identifies associations between `BRAF(A)` alterations and increased dependency with other downstream members of the MAPK signaling pathway including *MAP2K1*, *MAPK1*, *MITF*, and *DUSP4*. Associations with *MAP2K1*, *MAPK1*, and *MITF* are consistent with previous reports on conditional dependency on these genes in BRAF(V600E) melanoma [60, 61, 62].

**Figure 4:**
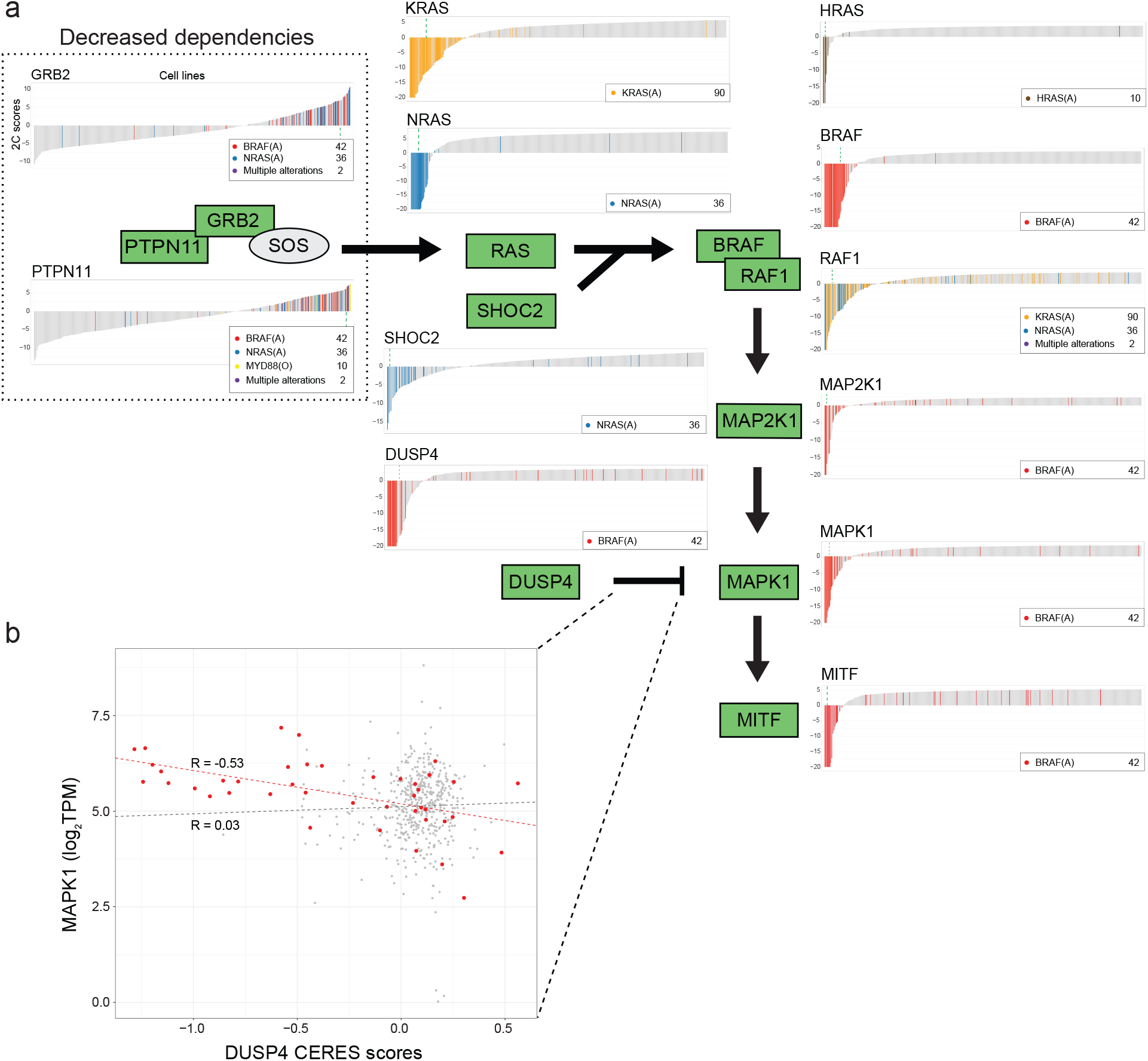
SuperDendrix identifies associations between genomic alterations and 2C differential dependencies in the MAPK pathway. (a) Activating mutations in *BRAF*, *KRAS*, *NRAS*, and *HRAS*, as well as other missense mutations in *MYD88* are approximately mutually exclusive and associated with 12 differential dependencies in the MAPK pathway (Same format as Fig. 2b). Alterations that activate RAS/RAF are associated with *increased* dependencies of 10 downstream genes in pathway. In contrast, these same alterations are associated with *decreased dependency* on 2 genes, *PTPN11* and *GRB2*, that are upstream activators of RAS. (b) CERES dependency scores of DUSP4 vs. expression of MAPK1. Cell lines with activating mutations in *BRAF* (red dots) show negative correlation between *DUSP4* dependency score and *MAPK1* expression (*R* = −0.53, *P <* 0.001), while no correlation is observed in cell lines without *BRAF* activating mutations (*R* = 0.03, *P* = 0.499).

The association between `BRAF(A)` alterations and increased dependency on *DUSP4* is intriguing because of conflicting reports on *DUSP4*’s role in cancer. *DUSP4* is reported to be a tumor suppressor that inhibits ERK1 and MAPK1 (ERK2) activity in the nucleus [63, 64]. As a tumor suppressor, *DUSP4* knockout would be expected to result in decreased dependency. However, there are also reports of high expression of *DUSP4* in colorectal cancers [63, 65] and skin cancers [66] with *RAS* or *RAF* alterations, suggesting that *DUSP4* may contribute to oncogenesis in these cancers. Our finding that cell lines with `BRAF(A)` have increased dependency on *DUSP4* is consistent with an oncogenic role. Since *DUSP4* is a negative regulator of *MAPK1*, we investigated the relationship between *DUSP4* dependency and *MAPK1*. We found that in cell lines with `BRAF(A)`, *DUSP4* dependency scores were significantly negatively correlated (*R* : −0.53*, P* ≤ 0.001; Pearson correlation) with expression of *MAPK1* (Fig. 4b); i.e. cell lines with `BRAF(A)` and highest MAPK expression were the most dependent on *DUSP4*. In contrast, in cell lines without `BRAF(A)`, there is no significant correlation between *DUSP4* dependency and MAPK expression (*R*: 0.03*, P* = 0.499). These observations are consistent with the Goldilocks principle described in [67], where precise levels of biological factors must be maintained for strong fitness, with either overdose or lack of oncogenic signal resulting in regression of tumor. In this case, *DUSP4* inhibition of *MAPK1* is most essential in cell lines with hyperactive MAPK signaling due to `BRAF(A)` alterations.

SuperDendrix also identifies two associations between sets of alterations and *decreased dependency* on *PTPN11* and *GRB2* in the MAPK pathway. Specifically, we identify decreased dependency on *PTPN11* in cell lines with `BRAF(A)`, `NRAS(A)`, or `MYD88(O)` alterations and decreased dependency on *GRB2* in cell lines with `BRAF(A)` or `NRAS(A)` alterations. The decreased dependency on *PTPN11* is consistent with a previous report that cell lines with constitutive RAS or RAF signaling were insensitive to suppression of *PTPN11* [68]. Another study reports that MYD88, similar to RAF or RAS, activates MAPK during cell transformations [69], potentially explaining the association between `MYD88(O)` alteration and *PTPN11* dependency exclusive of `BRAF(A)` and `NRAS(A)` alterations. Interestingly, 7 of the 10 `MYD88(O)` mutations occur in the TIR domain (*P* = 0.11; binomial test) which is reported to modulate MYD88 activity via interaction with Toll and IL-1 receptors [70] (Fig. S6). While we are not aware of previous reports of associations with *GRB2*, it is intriguing that both proteins with decreased dependencies – PTPN11 and GRB2 – are *upstream* of the RAS/RAF mutations that result in constitutive MAPK signaling. Thus, it makes sense that cell lines with constitutive activation of RAS or RAF signaling are insensitive to upstream activators of RAS signaling, analogous with the insensitivity of RB1-deficient cell lines to knockout of upstream regulators *CDK6* and *CCND1* reported above (Fig. 3).

The remaining 16 associations beyond those in the three pathways described above include associations between members of the same protein complex, and associations in other cancer-implicated pathways. Associations in protein complexes include: increased dependency on *ARID1B* in cell lines with `ARID1A(I)` alteration [48], increased dependency on *SMARCA2* in cell lines with `SMARCA4(I)` alteration [71], and increased dependency on *STAG1* in cell lines with `STAG2(I)` alteration [72]. Notable associations in pathways include 2 associations in the Wnt pathway: increased dependency on *CTNNB1* in cell lines with `APC(I)` or `CTNNB1(A)` alterations and increased dependency on *TCF7L2* in cell lines with `APC(I)` or `CTNNB1(A)` alterations (see “Associations in Wnt pathway” in Supplementary Information).

We found that 22 of the 32 associations identified by SuperDendrix validated in the Score dataset of CRISPR screens from Behan et al. [16], where we consider an association to be validated if there is a significant difference (*P* ≤ 0.05; Wilcoxon rank sum test) in dependency scores between cell lines containing associated mutations and those without such mutations (Table S4). Many of the associations that did not validate were in cancer types with few cell lines in the Score dataset. For example, several associations with `BRAF(A)` did not validate in the Score dataset; however, this is not surprising since the majority of `BRAF(A)` mutations in the Avana dataset are in the 34 skin cancer cell lines, while the Score dataset contains only 4 skin cancer cell lines (Table S5). Further details are in sections “Comparison to other perturbation screen results” and “Validation on the Sanger CRISPR-Cas9 screen data” in Supplementary Information.

Finally, we compared the associations between mutations and dependencies identified by SuperDendrix to those identified by Tsherniak et al. [15] in RNAi data and by Behan et al. [16] in a different CRISPR dataset. SuperDendrix finds significant associations for 6.5% of differential dependencies. This is higher than 1.3% in Tsherniak et al. and 1.6% in Behan et al.

### Cancer-type-specific differential dependencies

Next, we investigated associations between differential dependencies and cancer type. We augmented the alteration matrix with 31 cancer-type features, each feature representing one of the 31 cancer types in the Avana dataset. We ran SuperDendrix on the augmented alteration matrix and identified 135 differential dependencies that are significantly associated (FDR ≤ 0.2) with alterations and/or cancer types (Table S6). These include the 32 differential dependencies identified above using SuperDendrix on alterations alone. 18 associations are identical to those found by SuperDendrix when run on alterations alone while the remaining 117 include at least one cancer type feature. Among these are 14/32 profiles that were identified in the SuperDendrix analysis of genomic alterations described above but with associated sets that include cancer types and result in higher SuperDendrix weights. For example, *MITF* dependency and `BRAF(A)` (SuperDendrix weight = 0.21) has stronger association with `Skin cancer` (SuperDendrix weight = −0.75).

Of the 117 associations that include a cancer type feature, 92 are associations with increased dependency upon gene knockout, with the remaining 25 associated with decreased dependency. These 117 differential dependencies are enriched (FDR ≤ 0.05) for 85 GO molecular function terms (Table S7). The most significant function in these terms is *regulatory region nucleic acid binding*: in particular, 41 of the 117 are annotated as human transcription factors in [73] (fold enrichment = 4.07, *P* ≤ 0.001), a greater enrichment than the 70 transcription factors found among all 492 differential dependencies (fold enrichment = 1.65, *P* ≤ 0.001). This enrichment of transcription factor dependencies is consistent with previous reports; e.g., [15] identified 49 transcription factors with strong lineage-specific dependencies from RNAi screens. Our results include 15 of these 49 as well as 26 additional transcription factor dependencies that were not reported in [15].

The 41 transcription factor dependencies with cancer-type-specific associations cluster into a number of interesting groups (Fig. 5). These include dependencies on *ISL1*, *HAND2*, and *MYCN* in neuroblastoma, all of which were recently reported as part of the core regulatory circuitry (CRC) in neuroblastoma and associated with superenhancers [74]. Other large classes of cancer-type dependencies are in skin cancer (8 dependencies), breast cancer (5), leukemia or lymphoma (13), and multiple myeloma (10).

**Figure 5:**
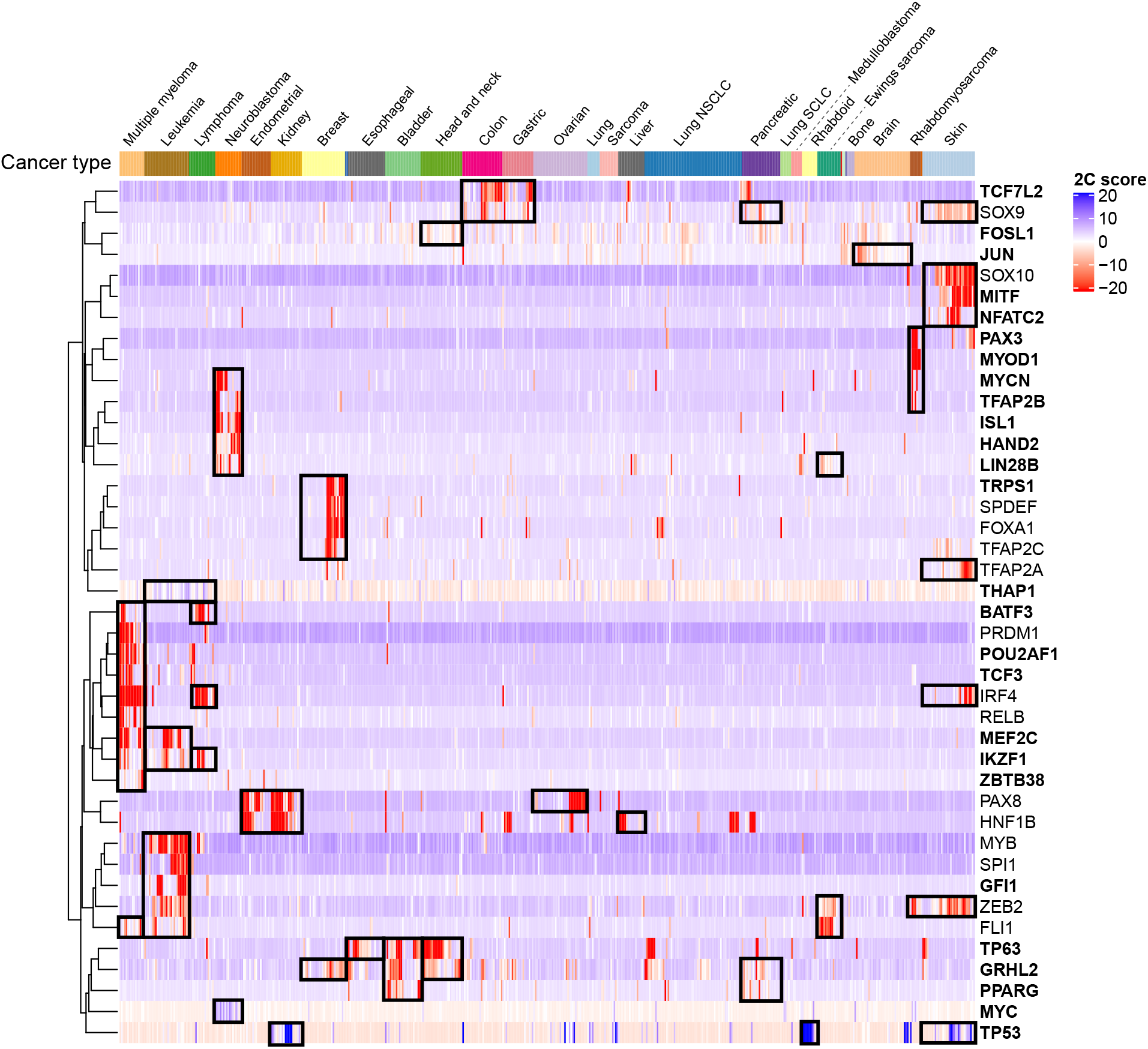
Transcription factors are the most common class of cancer-type-specific differential dependencies. Heatmap of 2C scores for 41 transcription factors that SuperDendrix identifies as cancer-type-specific differential dependencies. Dependency profiles are clustered within and across cancer types, with black boxes highlighting groups of prominent dependencies across cancer types. Bold text indicates transcription factors that were not reported in RNAi analysis [15]. Labels are shown for cancer types with at least 5 cell lines.

Two prominent cancer-type-specific associations identified by SuperDendrix are in the ID3-TCF3-CCND3 pathway in blood cancers (Fig. 6). Specifically, SuperDendrix finds an association between increased dependency on the transcription factor *TCF3* and the mutually exclusive set {`multiple myeloma, ID3(I), MEF2B(A)`}, as well as an association between increased dependency on *CCND3* and a mutually exclusive set {`leukemia, lymphoma`} (Fig. 6a). Chromosomal translocations resulting in *TCF3* fusion genes occur in various leukemias and lymphomas with differing reports regarding the importance of activation vs. inactivation of TCF3 in oncogenesis [75, 76]. The association between increased dependency on *TCF3* and `ID3(I)` or `MEF2B(A)` alterations is consistent with reports that ID3 inhibits TCF3 [77], and that TCF3 binds upstream of *MEF2B* [78, 79]. Thus, this association conforms to the model of oncogenic pathway addiction, with increased dependency on *TCF3* in cell lines with inactivating mutations in the *TCF3*-inhibitor *ID3* (analogous to the associations described above with the inhibitors *KEAP1* and *RB1*) or in cell lines with activating mutations in the downstream transcriptional activator *MEF2B*. *TCF3* has previously been suggested as a dependency in multiple myeloma cell lines, promoting tumorigenesis in cooperation with MYC [80]. In addition, the ID3-TCF3-CCND3 pathway is reported to be frequently perturbed in B-cell lymphoma with elevated MYC expression due to a chromosomal translocation [77]. Interestingly, the 5 cell lines with `ID3(I)` or `MEF2B(A)` mutations are 5 of the 7 B-cell lymphomas in the data set; moreover, the two `MEF2B(A)` alterations in these cell lines are D83V amino acid changes that are reported to be relevant in development of lymphoma [81]. Consistent with these reports, we found that the 5 B-cell lymphoma cell lines with increased dependency on *TCF3* and `ID3(I)` or `MEF2B(A)` alterations have high MYC expression (Fig. 6b). Interestingly, we find that the increased dependencies on *TCF3* and *CCND3* are approximately mutually exclusive, and the cell lines with both dependencies are the subset of B-cell lymphoma cell lines containing either `ID3(I)` or `MEF2B(A)` mutations (Fig.S7). Taken together, these results show increased dependency on the ID3-TCF3-CCND3 pathway in multiple myeloma, leukemia, and lymphoma, with the association in B-cell lymphomas modulated through alterations in *ID3* or *MEF2B*.

**Figure 6:**
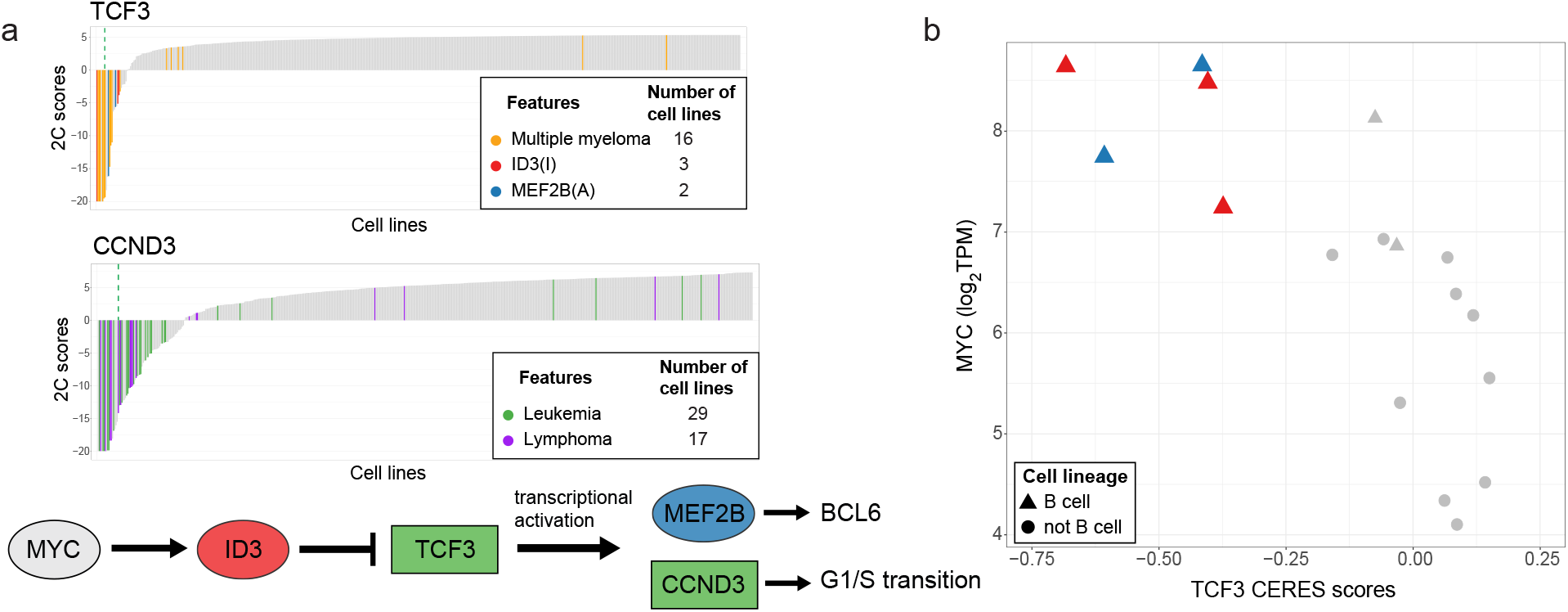
*TCF3* and *CCND3* dependencies in blood cancers. (a) (Top) SuperDendrix identified cancer-type-specific associations between increased dependency on *TCF3* and a mutually exclusive set {`multiple myeloma`, `ID3(I)`, `MEF2B(A)`} and also between increased dependency on *CCND3* and a mutually exclusive set {`leukemia, lymphoma`}. (Bottom) The ID3-TCF3-CCND3 pathway (Same format as Fig. 2b). (b) TCF3 CERES scores in lymphoma cell lines vs. expression of MYC. The 5 cell lines with *ID3* or *MEF2B* alterations are among cell lines with lowest *TCF3* CERES scores and highest MYC expression, and include 5 of 7 B-cell lymphoma cell lines in dataset.

The other 76 cancer-type-specific differential dependencies are enriched (FDR ≤ 0.05) for 46 GO molecular function terms (Table S8), with the top 3 enriched terms being *enzyme binding* (GO:0019899), *catalytic activity* (GO:0003824), and *kinase activity* (GO:0016301). These 76 differential dependencies include genes known to be overexpressed or predictive of prognosis for the associated cancer type such as increased dependencies of *MET* in brain cancer [82] and *LDB1* and *LMO2* dependencies in leukemia [83, 84]. Several additional associations correspond to dependencies on upstream regulators of cancer genes such as *MDM2* in skin and kidney cancers and *EGFR* in head and neck cancer.

A prominent grouping of cancer-type-specific differential dependencies are 6 genes in the IGF1R and PI3K pathways (Fig. 7a) across several cancer types. In the IGF1R pathway, we find increased dependency on *IGF2BP1*, *IGF1R*, *IRS1* or *IRS2* in neuroblastoma, Ewing’s sarcoma, multiple myleoma, and rhabdomyosarcoma. These dependencies are consistent with previous reports of dependencies on *IGF1R* in Ewing’s sarcoma and rhabdomyosarcoma [85]. SuperDendrix also find increased dependencies on *PIK3CA* and *BCL2* in some of these same cancer types, including *PIK3CA* in multiple myeloma and *BCL2* in leukemia, multiple myeloma, and neuroblastoma [86, 87]. These findings are consistent with IRS1/2’s role in activating the PI3K pathway [88]. Since dysregulation of the PI3K pathway results in tumor proliferation [89], all of these increased dependencies are consistent with a phenotype of oncogenic pathway addiction in the IGF1R/PI3K pathway.

**Figure 7:**
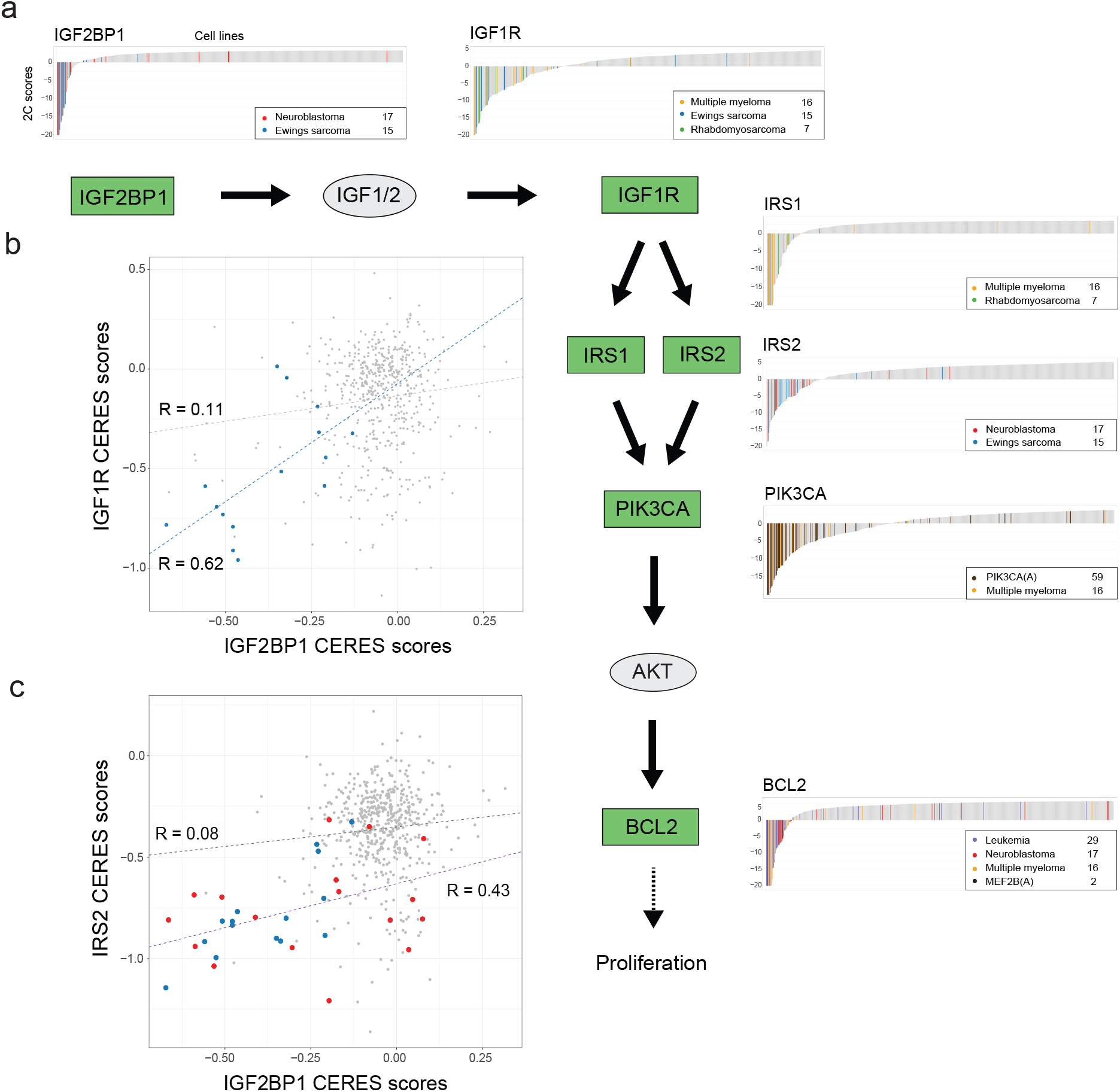
Cancer-type-specific differential dependencies in the IGF1R/PI3K pathway. (a) SuperDendrix identifies cancer-type-specific dependencies between 6 genes in IGF1R/PI3K pathway across multiple cancer types (Same format as Fig. 2b). (b) CERES scores of *IGF2BP1* and *IGF1R* are positively correlated (*R* = 0.62) in Ewing’s sarcoma cell lines (blue points), but only weakly correlated (*R* = 0.11) across other cancer types (gray points). (c) CERES scores of *IGF2BP1* and *IRS2* are positively correlated (*R* = 0.43) in Ewing’s sarcoma (blue points) and neuroblastoma (red points) cell lines, but only weakly correlated (*R* = 0.08) across other cancer types (gray points).

Since neuroblastoma, Ewing’s sarcoma, multiple myleoma, and rhabdomyosarcoma cell lines each exhibited dependencies on more than one gene in the IGF1R pathway, we examined whether individual cell lines harbored multiple dependencies. We found strong cancer-type-specific correlations between dependency scores of pairs of genes in the IGF1R pathway. These include correlations between *IGF2BP1* and *IGF1R* (*R* = 0.62*, P <* 0.001) in Ewing’s sarcoma (Fig. 7b) and between *IGF2BP1* and *IRS2* dependencies (*R* = 0.43*, P <* 0.001) in Ewing’s sarcoma and neuroblastoma (Fig. 7c). Importantly, these correlations are weaker in other cancer types (*R* = 0.11 and *R* = 0.08, respectively), and consequently were not reported in two recent studies [22, 90] that examined correlations between dependency profiles across all cell lines in DepMap. In addition, many of the cell lines with these correlated dependencies have CERES scores larger than the 0.6 threshold used to define dependency in DepMap [39]. Thus, the identification of these correlations relied on both SuperDendrix’s 2C scores and SuperDendrix’s ability to identify cancer-type-specific associations. At the same time, we find strong correlations between *IGF1R* with *IRS1* and *IGF1R* with *IRS2* across all cell lines, as previously reported in [22, 90]. We also find cancer-type-specific correlations between *CHUK* and *IKBKB* dependencies in lymphoma and multiple myeloma (Fig. S8).

Lastly, we found that 72 of the 117 associations identified by SuperDendrix are validated in the Score dataset [16], using the same test as described above (Table S6). Also as above, many of the associations that did not validate were in cancer types that were not well represented in the Score dataset including multiple myeloma, skin, and rhabdomyosarcoma (Table S5). Further details are in “Validation on the Sanger CRISPR-Cas9 screen data” in Supplementary Information.

## Discussion

We introduced SuperDendrix, a method that incorporates a principled statistical model and a practically efficient combinatorial algorithm to identify associations between differential dependencies and genomic alterations or cancer types. SuperDendrix scores differential dependencies using a two-component mixture model and identifies mutually exclusive sets of features – including genomic alterations and/or cell types – that are associated with each differential dependency. Application of SuperDendrix to data from Project DepMap – the largest publicly dataset of CRISPR-Cas9 loss-of-function screens in hundreds of cancer cell lines – identified 492 differential dependencies and inferred associations between 135 (27.4%) of these dependencies and sets of genomic alterations and/or cancer types. Many of these associations group into well-known cancer pathways such as MAPK, RB1, and IGF1R. SuperDendrix finds a higher fraction of differential dependencies are associated with genomic alterations compared to previous analyses of RNAi and CRISPR screens [15, 16, 24]. This illustrates advantages of the SuperDendrix method, including more stringent selection of differential dependencies and the search for sets of associated biomarkers; in contrast, existing approaches have very permissive definitions of differential dependencies or restrict to evaluating single-biomarker associations.

Interestingly, we observe striking consistencies between the directionality of dependencies (increased vs. decreased), the type of interactions (activating vs. inhibitory), and the position of dependencies and genomic alterations in pathways. Oncogenic mutations in upstream pathway genes – such as activating alterations in an oncogene or inactivating alterations in an tumor suppressor – are associated with *increased* dependencies in genes that are downstream in the same pathway and that promote cancer; e.g. *NFE2L2* dependency in cell lines with inactivating mutations in *KEAP1* and *MAPK1* dependency in cell lines with activating mutations in *BRAF*). These results are consistent with the notion that cancer cells develop addiction to an oncogenic pathway during cancer progression [17, 18]. On the other hand, oncogenic alterations in downstream pathway genes – such as activating alterations in an oncogene or inactivating alterations in a tumor suppressor gene – are associated with *decreased* dependencies on upstream genes of the same pathway; e.g. cell lines with inactivating mutations in *RB1* show decreased dependency on *CDK6*. These results show the importance of considering pathway topology in the design of cancer therapeutic strategies; for example, a current strategy for treating tumors with activating mutations in undruggable oncogenes is to inactivate downstream genes [91]. At the same time, current annotations of interactions in pathways should be interpreted with care and potentially revised with knowledge gained from perturbation experiments. For example, *DUSP4* is noted as a tumor suppressor due to its role in inhibiting MAPK signaling; however, we find increased dependency on *DUSP4* in cell lines with activating mutations in *BRAF* suggesting that DUSP4 contributes to maintaining the balance of MAPK signaling in *BRAF* mutant tumors. These results suggest DUSP4 as a potential therapeutic target for cancer treatment. Our results also provide further predictions about the the functional consequences of individual non-synonymous mutations and the function of individual genes. For example, we find that previously unannotated mutations in the dimerization domain of *KEAP1* are associated with increased dependency on its downstream target, *NFE2L2*.

SuperDendrix also identifies associations between differential dependencies and sets of cancer types or combinations of cancer types and genomic alterations. A large fraction (35%) of the cancer-type specific associations found by SuperDendrix involve increased dependencies on lineage-specific transcription factors. Many of these lineage-specific transcription factors have been previously reported to be highly expressed or correlated with poor prognosis in cancers of corresponding types. We also identify associations that include both cancer types and genomic alterations. For example, we find that increased dependency on *TCF3* in multiple myeloma cell lines or in cell lines with *ID3* or *MEF2B* alterations. These genomic alterations occur in the majority of B-cell lymphoma cell lines in the dataset. Another prominent cancer-type-specific association found by SuperDendrix is increased dependency on *IGF2BP1*, a regulator of insulin growth factor receptor *IGF1R*, in Ewing’s sarcoma and neuroblastoma. We anticipate that with larger cohorts, there will be increased opportunities to identify these more subtle associations that include both cancer types and genomic alterations.

There are a number of directions for future work, both the analysis and in further development of SuperDendrix. First, our current analysis does not include copy number aberrations (CNA) and DNA methylation changes, both of which are likely to be associated with differential dependencies. Since these alterations span larger genomic distances than single-nucleotide mutations, they require more careful decomposition into specific alteration events [92]. Since CNA occur frequently in solid tumors, we may be underestimating the number of associations in these cancer types. Second, alternative dependency scores could be used as input to SuperDendrix; recent studies have demonstrated that CERES scores can be affected by other covariates and confounding variables [93, 94]. Third, we found that some differential dependencies are associated with multiple sets of features (e.g. increased dependency on *TCF7L2* and the sets {`APC(I)`, `CTNNB1(A)`} and {Colon, Gastric}). Extending SuperDendrix to simultaneously identify multiple sets of features might identify additional such dependencies, as previously shown for multiple sets of mutually exclusive mutations [95]. Finally, SuperDendrix is a general algorithm that can be used to find associations between binary features (e.g. germline or somatic mutations, cell types) and quantitative phenotypes (e.g. drug response, cell size). It would be interesting to analyze these other phenotypes using SuperDendrix, particularly drug response data from The Genomics of Drug Sensitivity in Cancer (GDSC) database [96], and compare against other methods [97, 98] that have been designed specifically to identify associations between drug response and genomic features.

## Methods

### SuperDendrix algorithm

We introduce a new algorithm, SuperDendrix, to identify sets of binary features (e.g. genomic alterations or cell types) that are approximately mutually exclusive and associated with a continuous-valued phenotype. The inputs to SuperDendrix are:

1. An *l × n* matrix *P* = [*p_gj_*] of *l* quantitative phenotypes measured in *n* samples. Each entry *p_gj_* is the score of phenotype *g* in sample *j*. Each row of the phenotype matrix corresponds to a phenotype profile.
2. A list of binary features (e.g. somatic alterations) for each sample.
3. (Optional) Categorical information (e.g. cell type) of each sample.

While SuperDendrix is a general-purpose algorithm, here we describe the specific application where the phenotype scores are dependency scores from gene perturbation experiments and the binary features are genomic alterations (and optionally cell types). SuperDendrix includes three modules: (1) A module to identify and score differential dependencies using a two-component mixture model and to select genomic and cell-type features using mutation annotations; (2) A module to find sets of alterations/features that are approximately mutually exclusive and associated with differential dependencies using a combinatorial optimization algorithm; (3) A module to perform model selection and to evaluate statistical significance of associations.

### Identifying differential dependencies and selecting genomic features

The first module in SuperDendrix includes two steps: the identification and scoring of differential dependencies and the selection of genomic and cell-type features. In the first step, we assume that a gene perturbation leads to two population of samples: a minority of samples are *responsive* to the perturbation while the remaining samples are unresponsive and have scores derived from a *background* distribution. Thus, we assume that the dependency scores are distributed according to a two-component mixture model. We fit each dependency profile with a *t*-distribution and with a mixture of two *t*-distributions, using the *t*-distribution to model high variance in the dependency scores [99]. We use the Bayesian information criterion (BIC) [100] to select between the one-component or two-component models; we refer to genes whose dependency profiles are better fit by a two-component mixture as *differential dependencies*.

For each differential dependency *g* and sample *j*, we compute the 2C score, or differential dependency score, 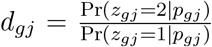, the log ratio of the posterior probabilities that the observed score is from component 2 or component 1. We compute posterior probabilities by fitting the dependency scores to a mixture of two *Gaussian distributions*. We choose component 1 to be the component with smaller mean so that negative 2C scores indicate decreased viability, or *increased dependency* in response to knockout. Conversely, positive 2C scores indicate *decreased dependency* in response to knockout. We assume that a minority of cell lines are responsive to gene knockout and thus refer to the component that contains fewer cell lines as the *responsive* component and the component with more cells lines as the *background* component. We define the 2C profile, or differential dependency profile, *d_g_* = (*d_g_*_1_*, …, d_gn_*) to be the differential dependency scores across all samples. Profile *d_g_* is an *increased dependency* if its responsive component contains cell lines with negative 2C scores (increased dependency) and a *decreased dependency* if its responsive component contains cell lines with positive 2C scores (decreased dependency).

In the second step, we construct a genomic alteration matrix *A* that includes annotated genomic alterations and (optionally) cell-type features. We construct *A* from non-synonymous somatic mutations in cancer genes in the OncoKB database [36]. We first classify each gene in OncoKB as **Oncogene**, **Tumor suppressor**, or **Other**based on “Oncogenicity” and “Alteration” columns in the OncoKB annotation as follows. For each gene, we select mutations with the following oncogenicity: “Likely Oncogenic,” “Oncogenic,” or “Inconclusive.” We classify genes that include “Deletion” or “Truncating Mutations” in the Alteration column as **Tumor suppressor**genes. We classify genes containing only “Inconclusive” mutations as **Other** genes. Finally, we classify each of the remaining genes as **Oncogene**. For each gene GENE in OncoKB, we group mutations from the input list into **A**ctivating (oncogene), **I**nactivating (tumor suppressor), or **O**ther alteration features which we label as GENE(A), GENE(I), and GENE(O) using the OncoKB annotation according to the following rules:

1. If GENE is **Oncogene**, its missense mutations that are included in the list of amino acid changes in OncoKB are grouped into a feature, GENE(A). Mutations in GENE that are not annotated in OncoKB are grouped into a feature, GENE(O).
2. If GENE is **Tumor suppressor**, its missense mutations that are included in the list of amino acid changes in OncoKB and mutations annotated as nonsense, nonstop, frameshift, start codon, stop codon, splice site, de novo start out of frame are grouped into a feature, GENE(I). Mutations in GENE that are not annotated in OncoKB are grouped into a feature, GENE(O).
3. If GENE is **Other**, then mutations are grouped into a feature, GENE(O).

Using the OncoKB alteration features derived above, we construct a genomic alteration matrix *A* = [*a_ij_*] of *m* OncoKB alteration features across *n* samples where *a_ij_* = 1 if alteration *i* occurs in sample *j* and *a_ij_* = 0 otherwise.

Next, we generate binary features that represent the cell type of each sample using information from metadata such as the primary tissue. In the application in this paper, we use cancer types as the cell-type features. Each cancer-type feature is marked as 1 for samples of that type, and as 0 for samples of other types. Note that the cancer-type features are mutually exclusive by definition. We now combine the two sets of features and create an augmented binary feature matrix *A* of *m* OncoKB alteration features and *q* cancer-type features across *n* samples.

### Finding feature sets associated with differential dependencies

The second module in SuperDendrix finds a subset *M\** of features (rows in *A*) that are: (i) most associated with differential dependency profile *d_g_*; and (ii) approximately mutually exclusive.

First, for each score *d_gj_* of differential dependency *g* in sample *j* from the profile *d_g_*, we define a normalized score 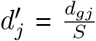 where 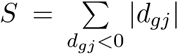 if *d_g_* is an increased dependency and 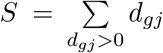 if *d_g_* is a decreased dependency. Then, we define a weight function *W* (*M*) that quantifies how well a subset *M* = (*m*_1_*, …, m_k_*) of features satisfies properties (i) and (ii). For the weight function *W* (*M*), we generalize the weight function defined in [31] to measure the mutual exclusivity between mutations. Specifically, for a set *M*, let Γ(*M*) be the subset of samples with alterations in *M*, *c_j_* (*M*) be the number of alterations in *M* that occur in sample *j*, and *ρ_j_* be a penalty term for alterations in *M* that co-occur in sample *j*. When searching for association to increased dependency, *ρ_j_* is equal to *−|d′_j_|*; when searching for association to decreased dependency, *ρ_j_* is equal to *|d′_j_|*. We define

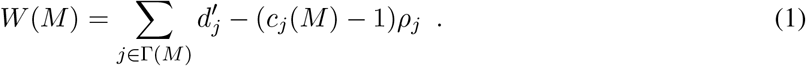

If the alterations in *M* are mutually exclusive then *c_j_* (*M*) = 1 for all *j* and thus *W* (*M*) is the sum of differential dependency scores for all altered samples. If *c_j_ >* 1, then sample *j* has alterations in more than one feature in *M*, and thus we penalize the weight *W* (*M*) for each additional alteration. Note that if the features that co-occur in a sample are GENE(I) and GENE(O) alterations, we do not penalize the weight. This is motivated by the two-hit hypothesis [101] which states that both alleles need to be mutated for gene inactivation. To see that the weight *W* (*M*) is a straightforward generalization of the Dendrix weight introduced in [31] we consider the following reformulation, in which Γ(*m*) denotes the set of samples with feature *m*.

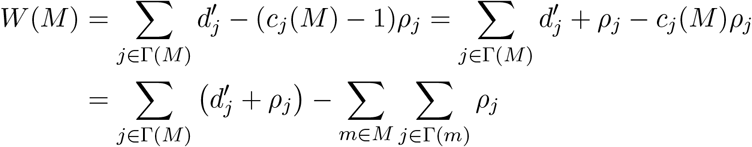

In the case where all samples have equal score, i.e., *d*′_*j*_ = 1, and *ρ*_*j*_ = |*d*′_*j*_| = 1 for all *j*, the supervised Dendrix weight *W* (*M*) simplifies to 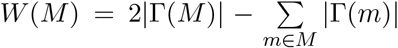, which is the original Dendrix weight defined in [31].

Following the nomenclature in machine learning, the problem considered in Dendrix [31] of finding a mutually exclusive set of alterations is an “unsupervised” feature selection problem, while the problem solved by SuperDendrix is a “supervised” feature selection problem where we aim to identify a set of mutually exclusive features that “explain” a phenotype.

Next, we aim to find a set *M\** with optimal weight *W* (*M\**), which we define as follows.

*Problem* 1 (Optimal Weight Exclusive Target Coverage Problem (OWXTC)). Given an alteration matrix *A* and a differential dependency profile *d*, find a subset *M\** of rows satisfying

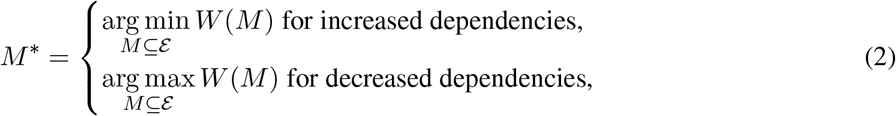

where *ε* is all subsets of rows in *A*.

OWXTC is NP-hard because it generalizes the Maximum Weight Submatrix Problem which was shown to be NP-hard in [31] for the special case where **d**′ = (1, 1*, …, * 1) and *ρ* = **d**′. We also define the cardinality-constrained version *k*-OWTXC of OWXTC in which *ε* is all subsets of size at most *k*.

We formulate the OWXTC as an integer linear program (ILP) as follows. First, we define binary variables *x_i_*, for each row 1 *≤ i ≤ m*, and *y_j_*, for each column 1 *≤ j ≤ n*, with the interpretation

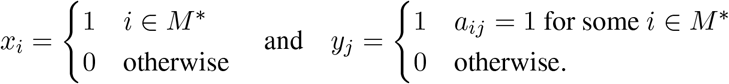

Then the OWXTC in the case of increased dependency is equivalent to the following ILP.

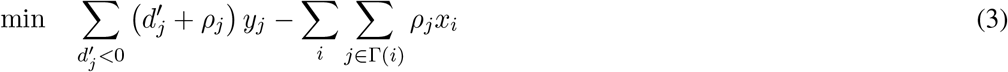

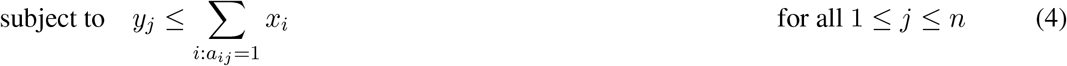

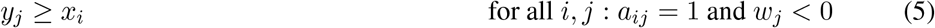

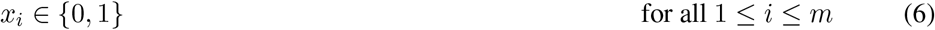

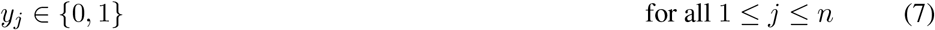

For finding associations with decreased dependencies, we replace min by max in Equation (3). For the cardinality-restricted version, we add the inequality

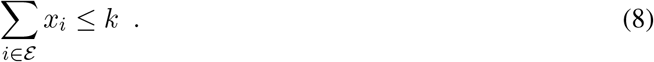

Note that the SuperDendrix weight and the ILP are similar, but not identical, to those presented in [26]. The differences are discussed in Supplementary Information.

### Evaluating statistical significance of associations

The third module of SuperDendrix consists of two steps. First, since the optimal size *k* = *M\** of the feature set is unknown, we perform model selection using a conditional permutation test to evaluate the contribution of each alteration to the weight *W* (*M\**). For each feature *m* in *M\**, we compare the weight *W* (*M\**) to the distribution of the weight 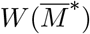, where 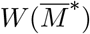 is the weight obtained when mutations of the feature *m* are permuted across samples. We compute the empirical *P*-value as 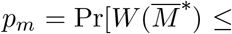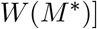 (increased dependency) or 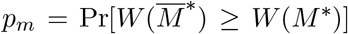 (decreased dependency) over 10,000 permutations and remove *m* with the largest *P*-value only if *p_m_ >* 0.0001. We repeat the above process until we obtain a feature set 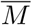 which only contains features with *p_m_* ≤ 0.0001.

Next, we evaluate the statistical significance of the association between feature set 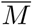 and differential dependency profile **d**by running SuperDendrix on random feature matrices 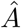 with fixed row and column sums (numbers of alterations per gene and sample, respectively) [102]. Note that we generate these random matrices using *all* alterations (i.e. including mutations not annotated in OncoKB), and then use the first module in SuperDendrix to select the OncoKB alteration features. We compare the weight 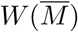 to the distribution of the weight 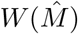, where 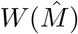 is the optimal weight computed from a random feature matrix 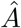. We compute the empirical *P*-value as 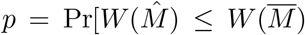 (increased dependency) or 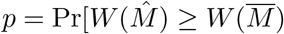 (decreased dependency) over 10,000 random feature matrices.

After computing *P*-values of the feature sets for each differential dependency, we compute false discovery rate (FDR) using Benjamini-Hochberg procedure [103] for multiple hypothesis correction.

#### Implementation and Software

We implement SuperDendrix using Python 3. We use the R package, EMMIXskew [104], to fit *t*-distribution mixture models to dependency scores. We use the Python scikit-learn library [105] to fit Gaussian mixture models to dependency scores and to compute the 2C scores. We solve the ILP using the Gurobi software [106]. SuperDendrix software and our experiments are available at at https://github.com/raphael-group/superdendrix.

### Bioinformatics and Data processing

We downloaded the Avana [19Q1/2.19.2019] dataset from the DepMap data portal^1^ [39]. This dataset contains dependency scores – computed using the CERES algorithm – for 17,634 CRISPR-Cas9 gene knockouts across 558 cancer cell lines. We normalize each of 17,634 dependency profiles by converting CERES scores to z-scores as described in [39] before applying SuperDendrix. After running the first module of SuperDendrix, we obtain 492 differential dependencies that are better fit by a mixture of two *t*-distributions; 412 increased dependencies and 79 decreased dependencies.

We downloaded mutation data [19Q1/2.19.2019] for the same cell lines from the Cancer Cell Line Encyclopedia (CCLE) [43] using the same DepMap data portal. This dataset includes mutation data for 554 of the 558 cell lines in the CRISPR-Cas9 dataset. We applied SuperDendrix to 788,985 non-synonymous mutations. After running the first module of SuperDendrix we obtain an alteration matrix containing 566 alteration features (49 GENE(A), 229 GENE(I), and 228 GENE(O) alteration features) in 355 genes in 554 cell lines. Note that these alteration features do not overlap with the list of recently identified “passenger hotspot” mutations caused by preferential APOBEC activity in DNA stem loops [107]. To derive cancer-type features, we used the “primary tissue” and ‘secondary tissue” columns in the DepMap cell line metadata^2^ and fixed annotation errors and merged rare cancer sub-types. Our annotation of cancer types is in “primary tissue” column in our curated cell line data (Table S9). We use this annotation to construct 31 binary cancer-type features representing the cancer types of DepMap cell lines where each feature has a value 1 for cell lines of that cancer type and 0 for other cell lines.

We run SuperDendrix using sets of at most 3 alterations and sets of at most 5 alterations and cancer types.

### Web browser for genetic dependency and mutation data

We release a public, open-source web browser to view and explore SuperDendrix results. Users can choose which genetic dependency profile and which mutations they want to view, or can preload an association identified as significant by the SuperDendrix software. The browser displays a waterfall plot, indicating the dependency score and mutation status in each cell line. It also includes two bar plots on top of the waterfall plot that indicate tissue type and number of mutations per cell line. Users can interact with the plots by scrolling over bars in the waterfall plot. On mouse over, the browser displays tooltips listing information about the given cell line such as tissue type. Users can also select a range of cell lines in the bar plot at the top to zoom in. The plots provide an easy way to quickly assess whether the dependency scores in cell lines with user-specified alterations or cancer types are extreme relative to the other cell lines. The code for the SuperDendrix browser is open-source at https://github.com/lrgr/superdendrix-explorer, and the browser itself is publicly available at https://superdendrix-explorer.lrgr.io/.

### Comparison to other methods

As noted in the introduction, there are two other methods to find associations between mutually exclusive alterations and gene perturbation scores: REVEALER [25] and UNCOVER [26]. REVEALER uses a greedy method to find mutually exclusive alterations associated with continuous phenotype. As noted in [26], the greedy method is slow and not scalable to the large-scale Avana dataset containing thousands of dependency profiles, and therefore was not compared with SuperDendrix. UNCOVER was developed concurrently with our development of SuperDendrix, and also solves a combinatorial optimization problem. However, there are several key differences between SuperDendrix and UNCOVER.

1. UNCOVER is applied directly to dependency scores, while SuperDendrix first identifies and scores differential dependencies using a mixture model.
2. UNCOVER combines all mutations in a gene into a single gene-level alteration, while SuperDendrix creates different alteration features (GENE(A), GENE(I), or GENE(O)) according to OncoKB annotations.
3. UNCOVER uses a different objective function in the optimization with positive and negative scores having asymmetric contribution to the objective.
4. UNCOVER lacks a model selection step, and does not control for variability in the number of mutations across cell lines during its statistical test.

Further details of these differences are in Supplementary Information.

We compared SuperDendrix and UNCOVER on the same Avana dataset, and found that the methodological differences between SuperDendrix and UNCOVER led to large qualitative and quantitative differences in results. We ran UNCOVER using CERES scores of 1,730 6*σ* profiles and 9,987 alteration features and 31 cancer-type features (see Supplementary Information for data processing) to search for a set of 3 associated alterations features and 5 alteration or cancer-type features. UNCOVER reported 139 sets of alterations containing a total of 417 alterations with significant association (FDR ≤ 0.20, Table S10), compared to 32 sets containing a total of 44 alterations for SuperDendrix. When the 31 cancer-type features were included, UNCOVER reported 722 associations compared to 135 for SuperDendrix (Table S11).

There are multiple reasons for the larger number of associations predicted by UNCOVER. First, 71 of the 139 significant associations identified by UNCOVER are associations between dependencies that are not identified as differential dependencies by SuperDendrix. These include dependencies on *ZNF592* and *MAP3K2* whose dependency score distributions are unimodal (Fig. S9). Second, UNCOVER does not include a model selection procedure, and thus always returns alteration sets of the requested size (3 and 5 in these experiments). Of the 24 differential dependencies where both UNCOVER and SuperDendrix reported associated alterations, UNCOVER’s associated sets included 72 gene-level alterations (24×3), while SuperDendrix sets contain a total of 34 alterations (including GENE(A), GENE(I), and GENE(O) alterations). 29 of the gene-level alterations identified by UNCOVER overlap the 34 alterations identified by SuperDendrix in the corresponding profile. The remaining 43 gene-level alterations found by UNCOVER are not included in SuperDendrix results. Notably, 33 of these 43 alterations contribute less than 20% to the corresponding weight of the alteration set. Across UNCOVER’s 139 total significant associations, 18% of the significant alterations 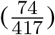 contribute less than 20% to the set’s weight. These alterations with small objective values are likely false positives. Finally, the permutation test used to evaluate statistical significance of UNCOVER’s results does not control for variability in the number of mutations across cell lines. We found that the number of significant associations reported by UNCOVER in a cell line is significantly correlated with the number of alterations in the cell line (Pearson correlation: *R* = 0.85 for alterations; and *R* = 0.83 for alteration and cancer-type features) (Fig. S10), indicating that some of the associations reported by UNCOVER are likely false positives. In comparison, the correlation is much weaker in SuperDendrix results (*R* = 0.54 for alterations only; and *R* = 0.27 with cancer types included).

## Acknowledgements

G.W.K. thanks the Fulbright Scholarship Program for support. This work is supported by US National Institutes of Health (NIH) grants R01HG007069 (B.J.R.) and U24CA211000 (B.J.R.).

## Supplementary Figures

**Figure S1:**
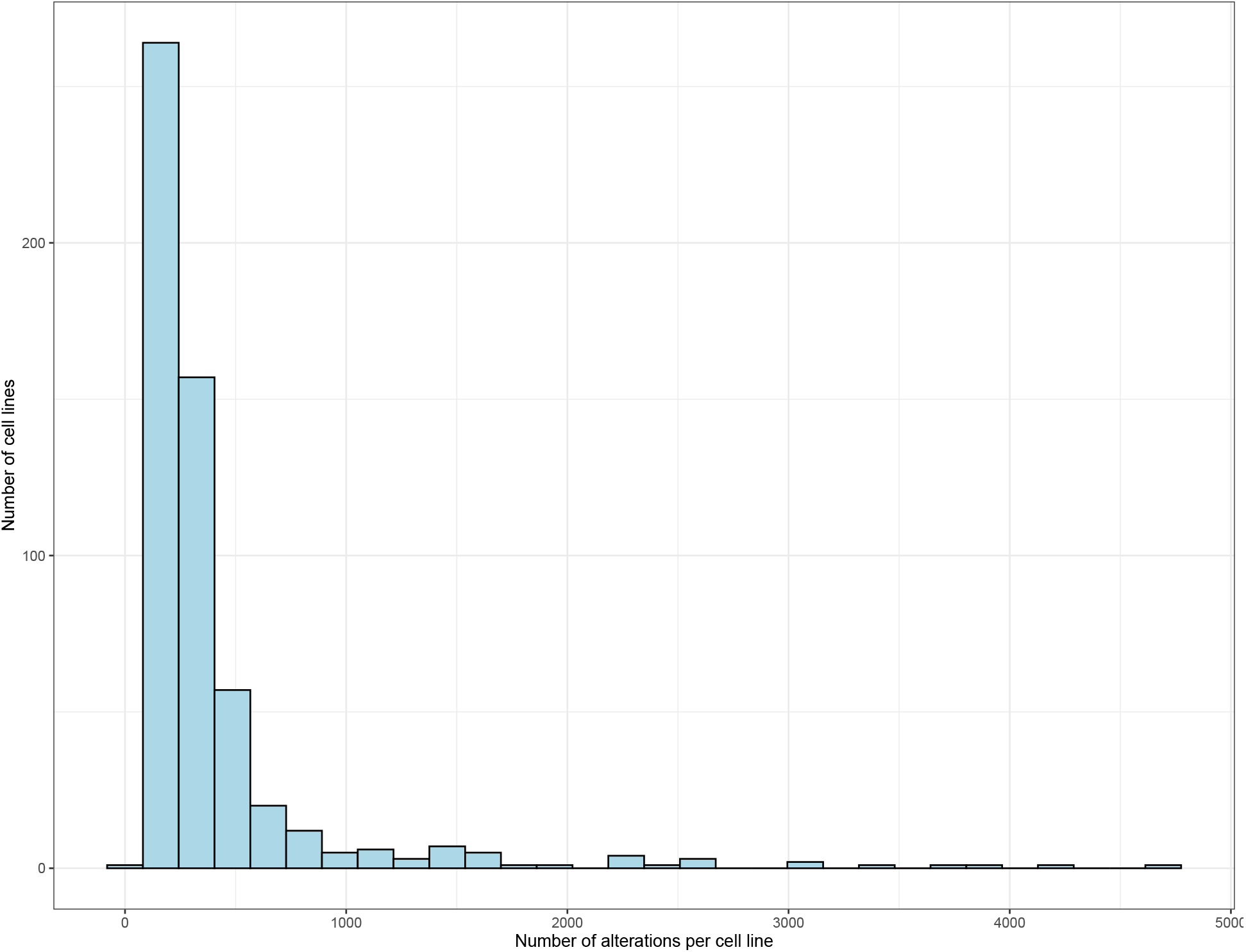
Number of genomic alterations in cell lines in DepMap. Distribution of all non-synonymous mutations in the cell lines from DepMap. Mean = 410.01 and standard deviation = 533.43.

Figure S2: **Results of fitting a mixture of two t-distributions to the 17,634 dependency profiles**. Attached as a supplement file.

**Figure S3:**
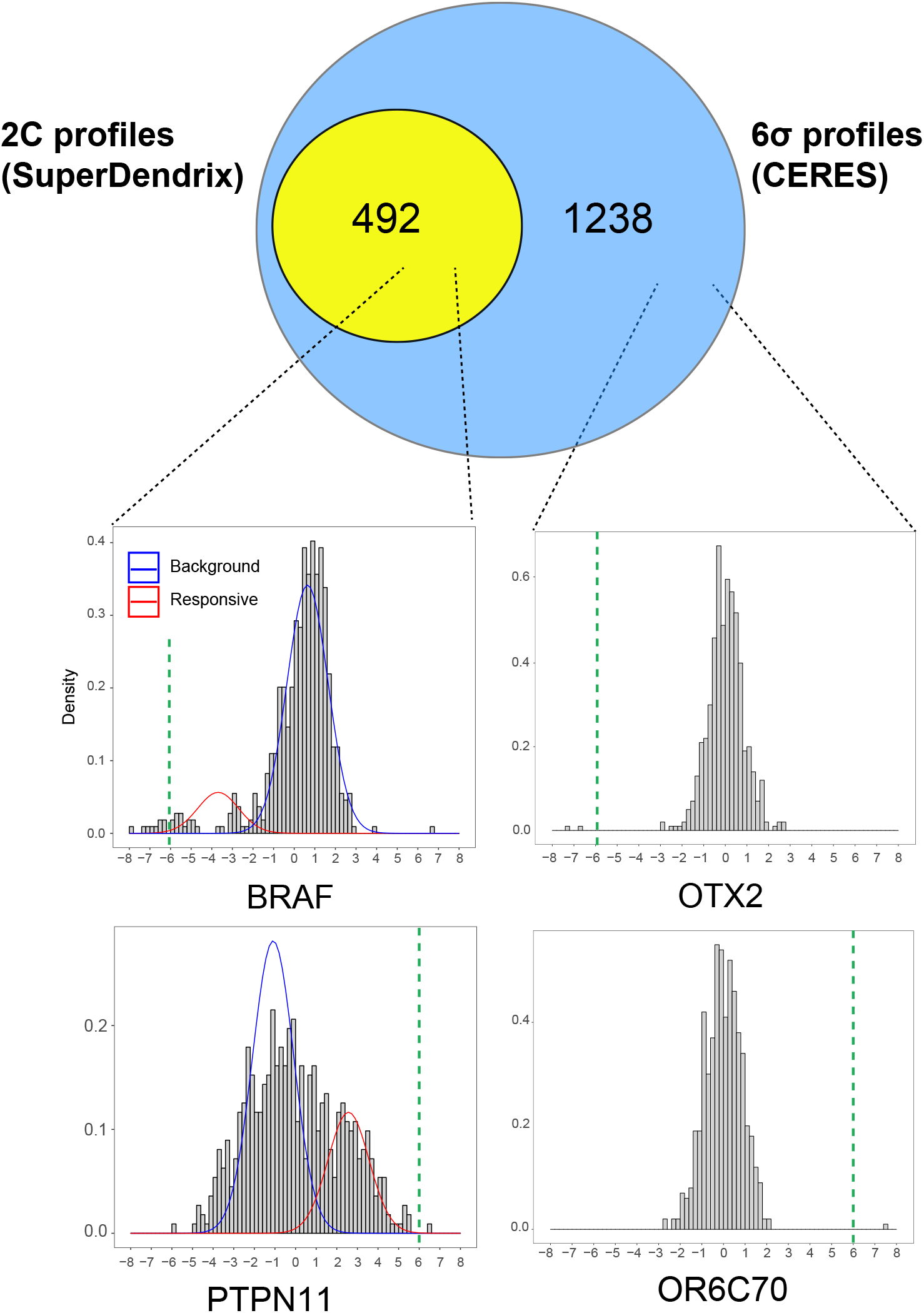
Comparison of differential dependencies found by SuperDendrix and CERES. (Top) Overlap between differential dependencies identified using the two component mixture model in SuperDendrix and the 6*σ* threshold for CERES z-scores as described in [15]. (Bottom) CERES z-score distributions for example profiles from each section of Venn diagram. The top row are profiles with increased dependency upon gene knockout, while the bottom row are profiles with decreased dependency. Red and blue curves indicate distributions in two component mixture, and green dashed lines represent 6*σ* thresholds for CERES z-scores.

**Figure S4:**
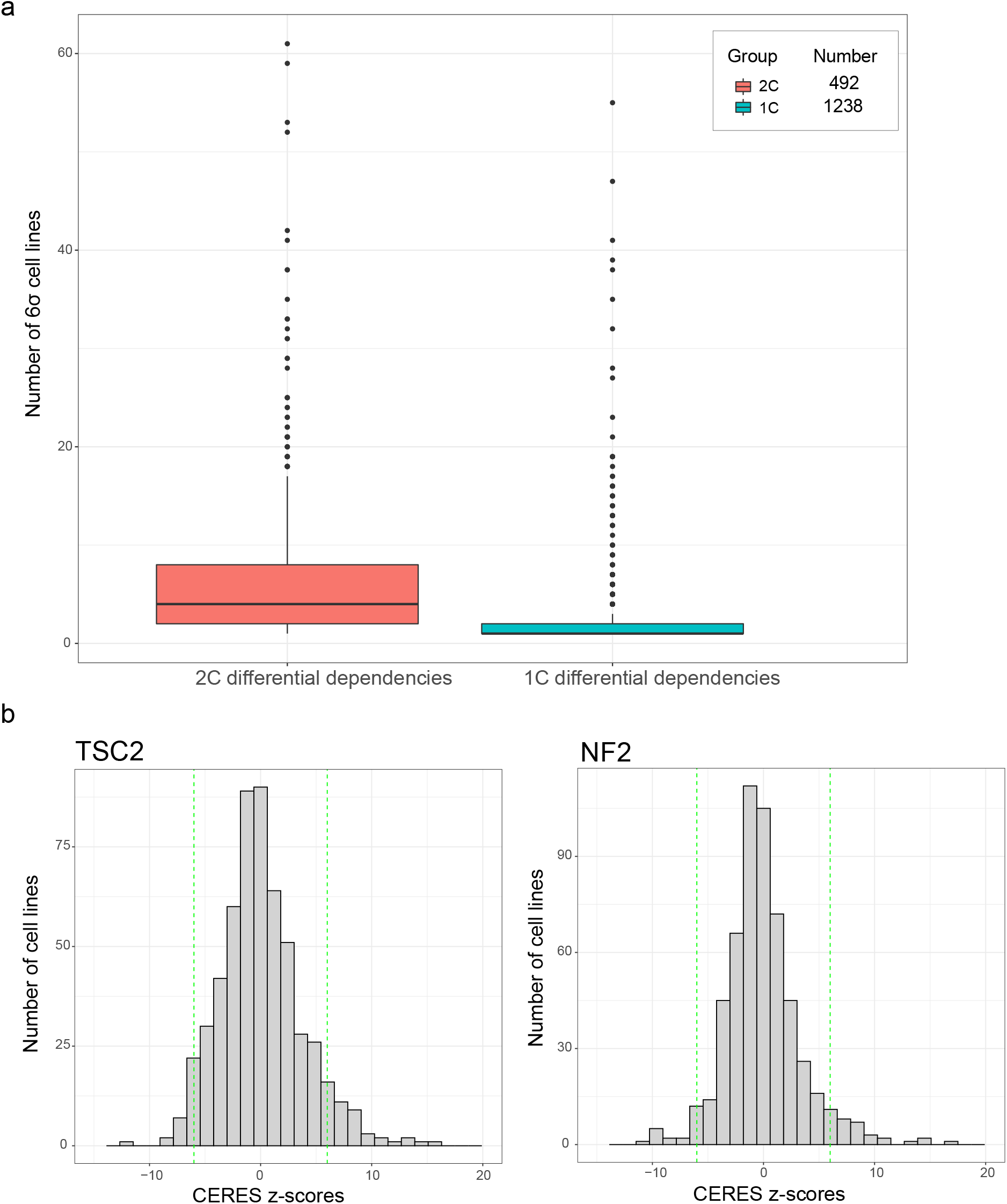
Number of 6*σ* outlier cell lines in 2C and 1C profiles. (a) Box plots of number of cell lines with CERES scores ≥ 6*σ* from the mean for 2C profiles and 1C profiles. 2C profiles contain higher number of 6*σ* cell lines than 1C profiles on average (Welch t-test, *P <* 0.001). (b) CERES z-score distributions of *TSC2* and *NF2*, the 1C profiles containing the highest number of 6*σ* outlier cell lines (55 and 47). The outlier distributions have wide standard deviations: 3.73 and 3.42. The green dashed lines represent 6*σ* thresholds for CERES z-scores.

**Figure S5:**
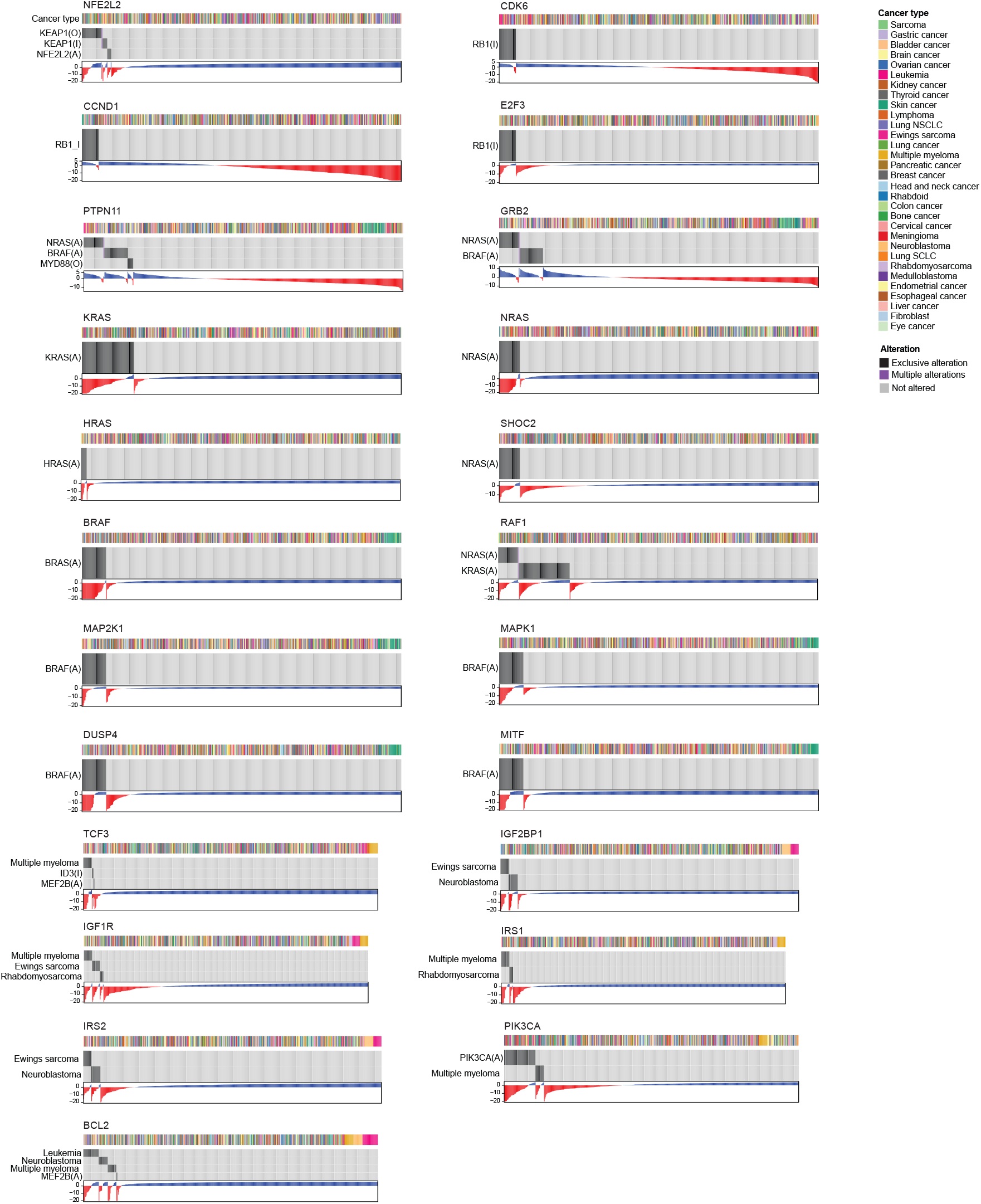
Alteration matrices and profiles for significant associations discovered by SuperDendrix. Blue boxes are cell lines that contain the corresponding alteration. Red boxes are cell lines that contain multiple alterations in each association.

**Figure S6:**
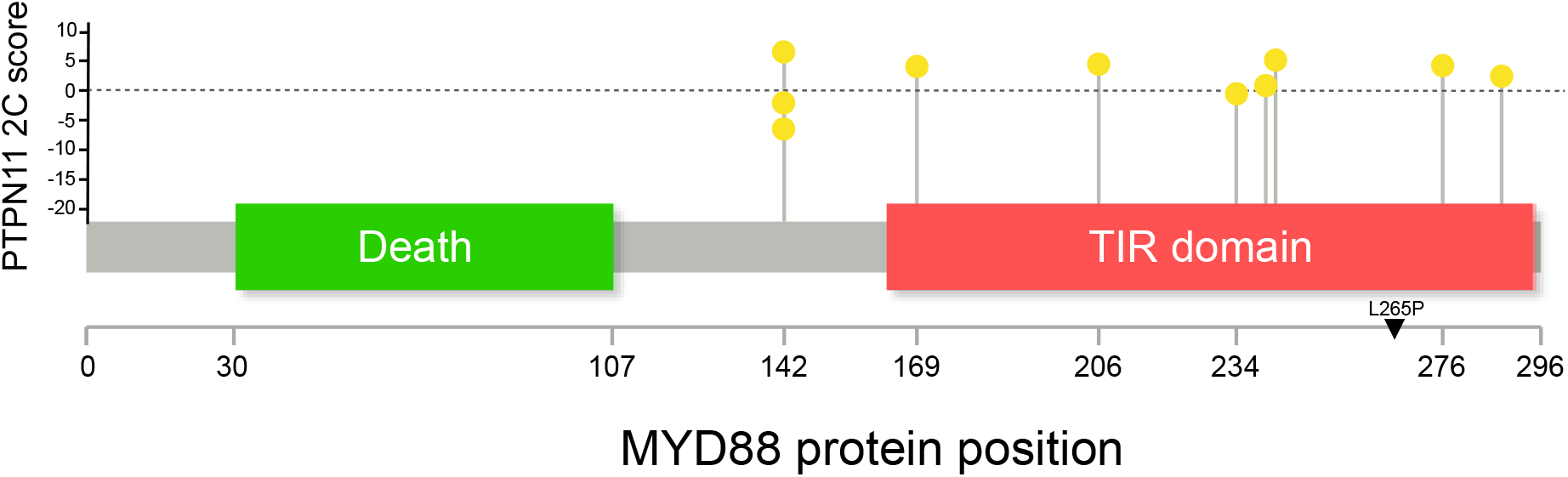
Locations of mutations in MYD88 protein that are annotated as Other. `MYD88(O)` mutations are associated with decreased dependency on *PTPN11*. 7 of the 10 `MYD88(O)` mutations occur in TIR domain (*P* = 0.11; binomial test). CERES scores for *PTPN11* are higher (decreased dependency) on average in the 7 cell lines with `MYD88(O)` mutations in TIR domain than the 3 cell lines with `MYD88(O)` mutations outside of TIR (*P* = 0.13; t-test).

**Figure S7:**
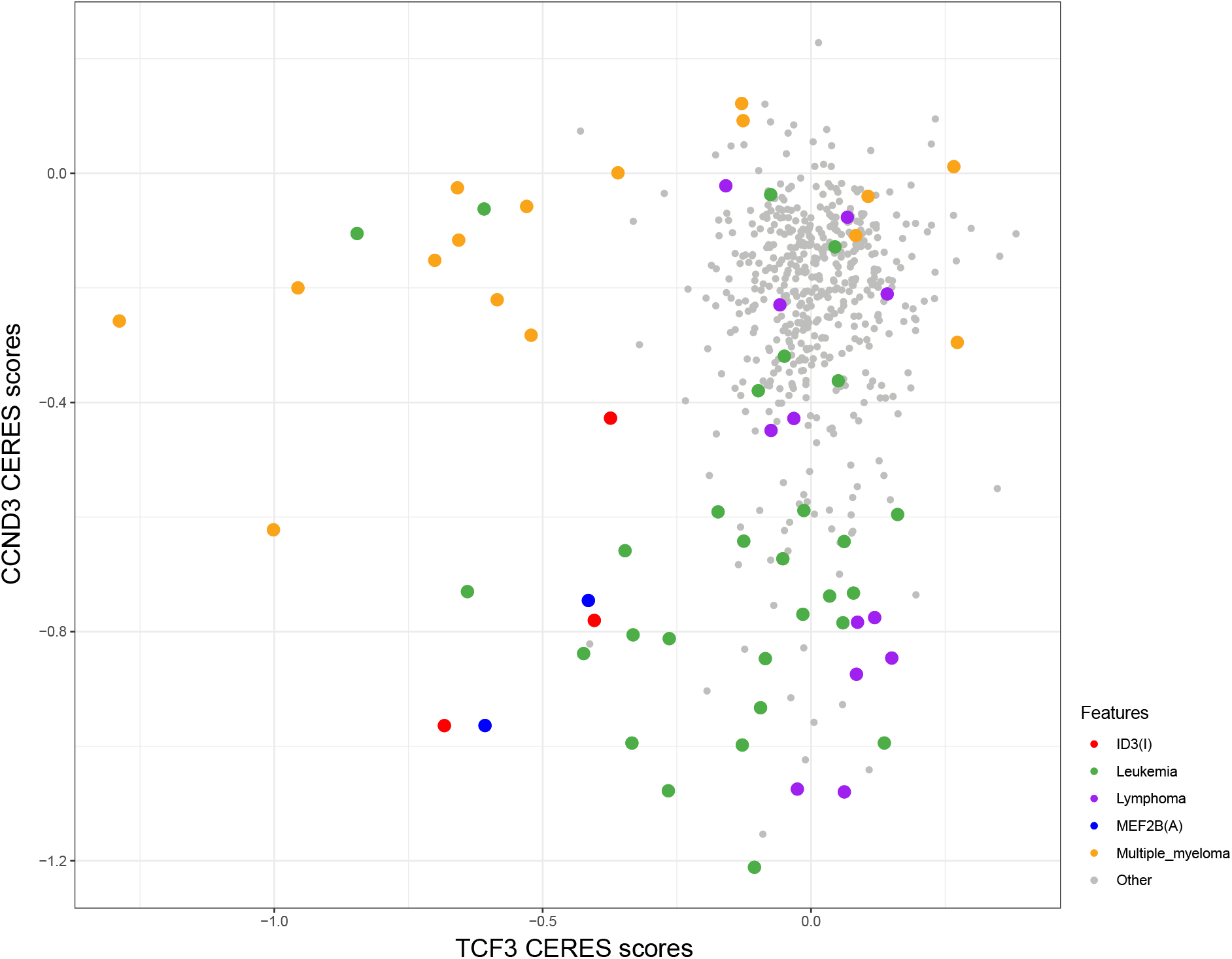
Increased dependencies on *TCF3* and *CCND3* in blood cancers. CERES dependency scores for *TCF3* (x-axis) and *CCND3* (y-axis) across cell lines with multiple myeloma (orange), leukemia (green), and lymphoma cell (purple) cell lines showing a mutually exclusive pattern with increased dependency on either *TCF3* and *CCND3*. In contrast, the subset of B-cell lymphomas containing `ID3(I)` (red) or `MEF2B(A)` (blue) mutations show increased dependencies on *both TCF3* and *CCND3*.

**Figure S8:**
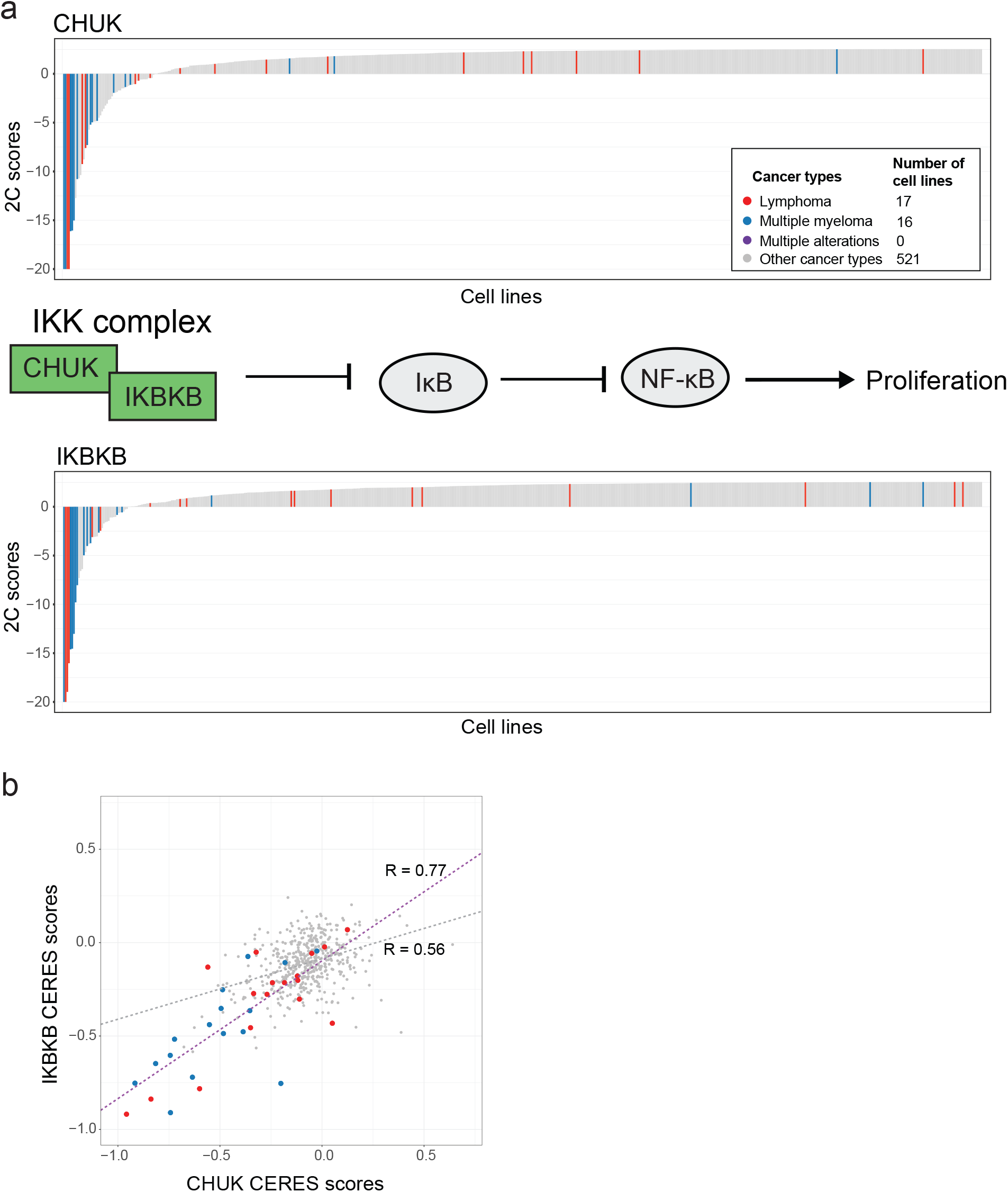
Cancer-type specific differential dependencies in NF-*κ*B pathway. (a) Lymphoma and multiple myeloma are associated with increased dependency on *CHUK* and *IKBKB*, the *α* and *β* subunits of IKK complex, that activate downstream NF-*κ*B pathway (Same format as Fig. 2b). This is consistent with previous reports of NF-*κ*B deregulation in blood cancers [108, 109, 110]. (b) CERES scores of cancer types that are differentially dependent in both *CHUK* and *IKBKB* profiles show positive correlation (*R* = 0.77, *P <* 0.001).

**Figure S9:**
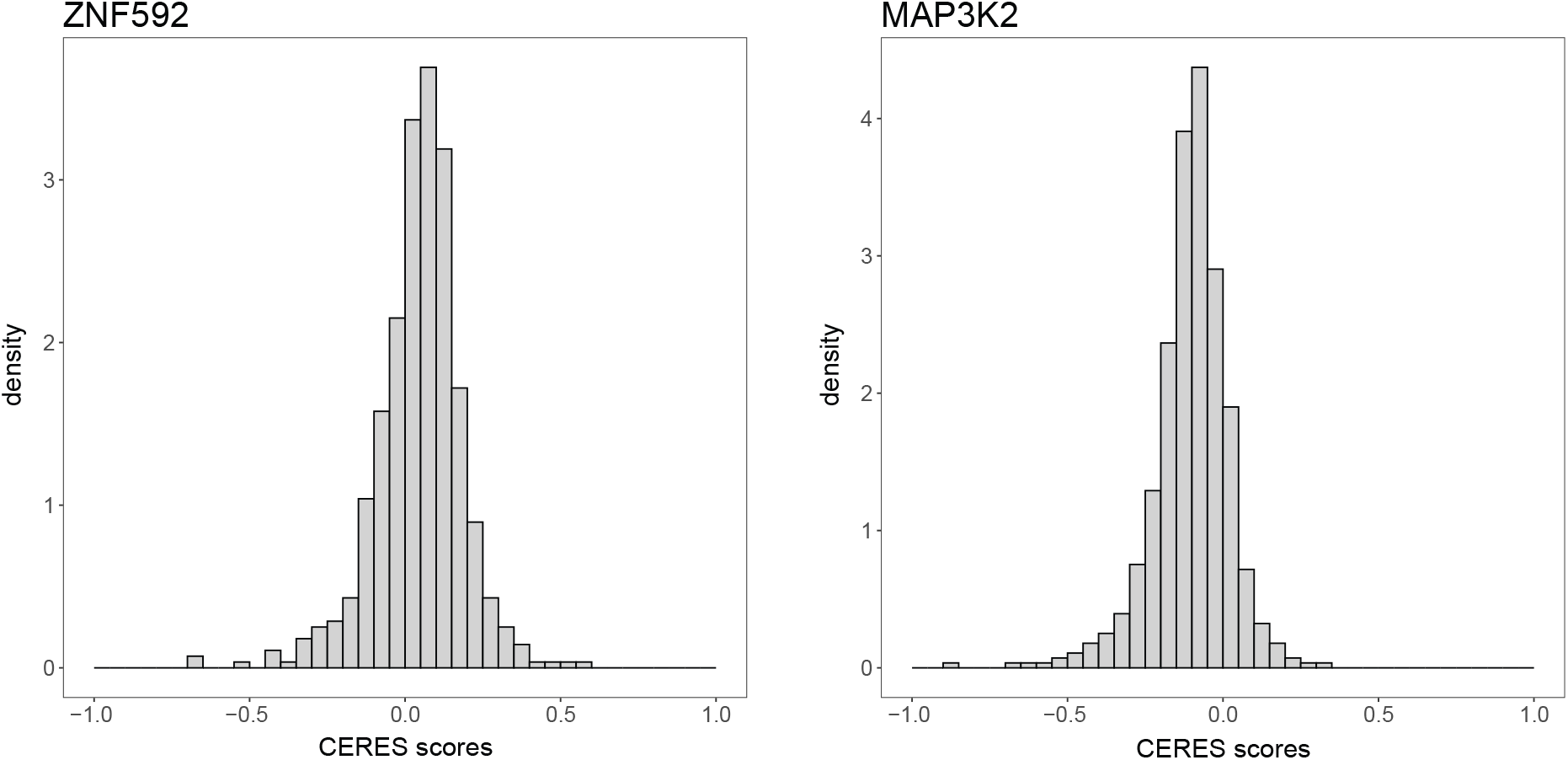
UNCOVER finds associations between alterations and dependencies with unimodal score distribution. CERES score distributions of *ZNF592* and *MAP3K2*. Each of these distributions is unimodal and better fit by a single t-distribution than by a mixture of 2 t-distributions.

**Figure S10:**
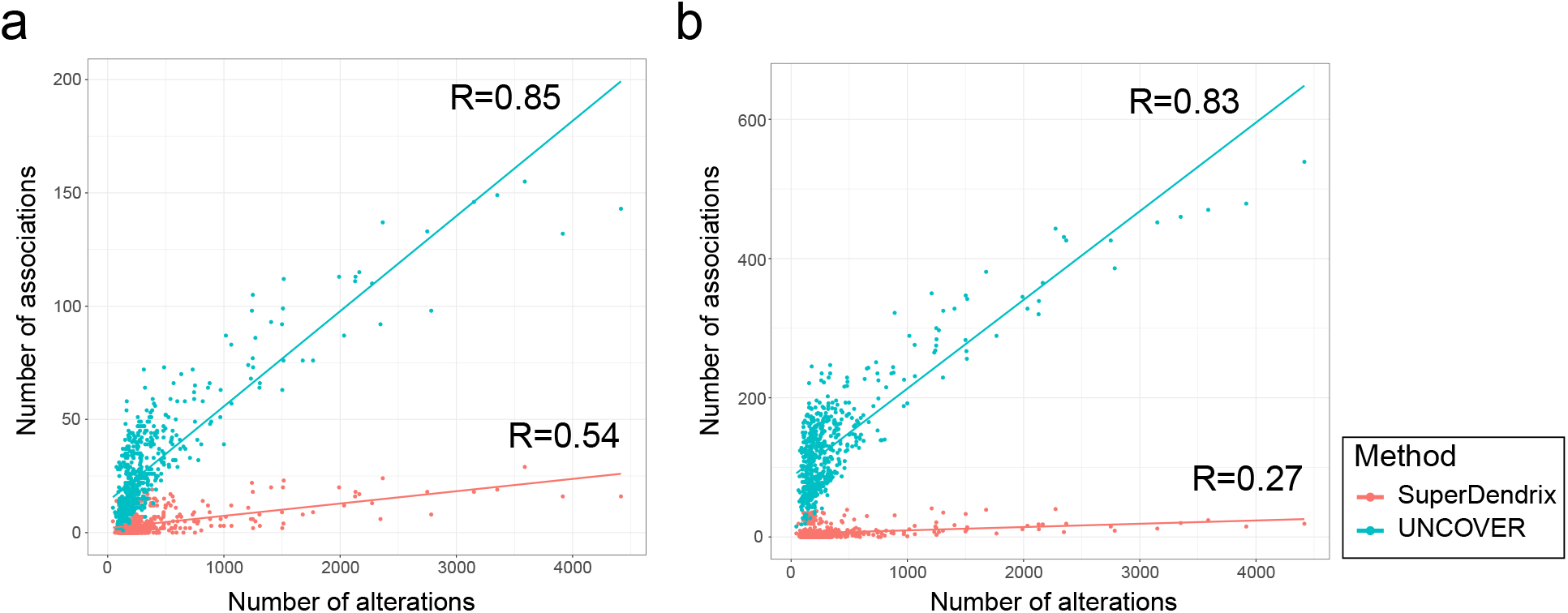
Correlation between total number of alterations and alterations from significant associations in each cell line. (a)-(b) Correlation was computed by comparing the total number of alterations (prior to OncoKB filtering) against the number of significant features in each cell line. (a) Correlation is lower in SuperDendrix for alteration results (SuperDendrix: *R* = 0.54, UNCOVER: *R* = 0.85). (b) Correlation is lower in SuperDendrix also for cancer type results (SuperDendrix: *R* = 0.27, UNCOVER: *R* = 0.83).

**Figure S11:**
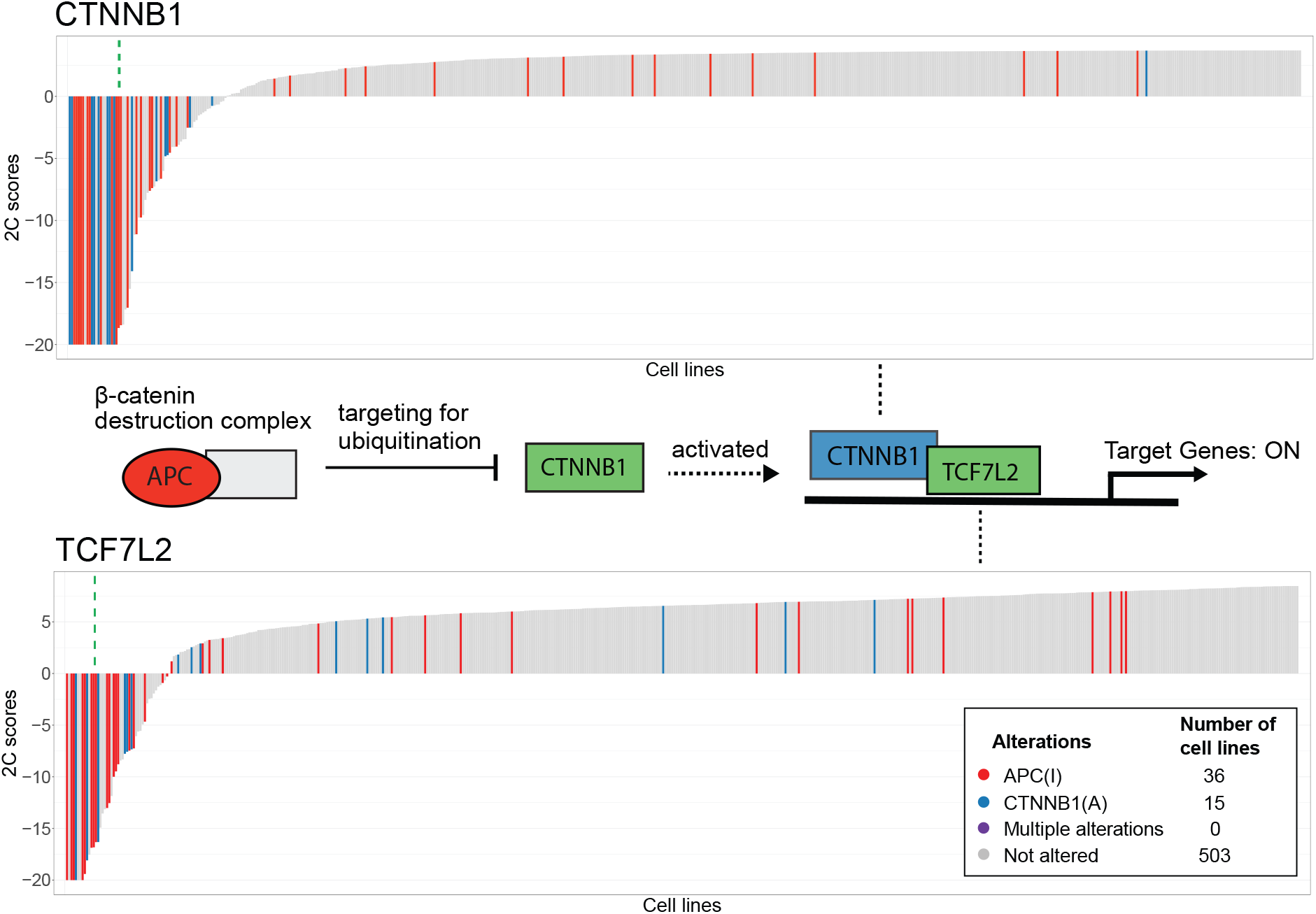
SuperDendrix identifies dependencies in the Wnt pathway. Cell lines with *APC* inactivating and *CTNNB1* activating alterations are associated with increased dependency on *CTNNB1* and *TCF7L2* (Same format as Fig. 2b). These associations are consistent with the known interactions on Wnt signaling pathway.

**Figure S12:**
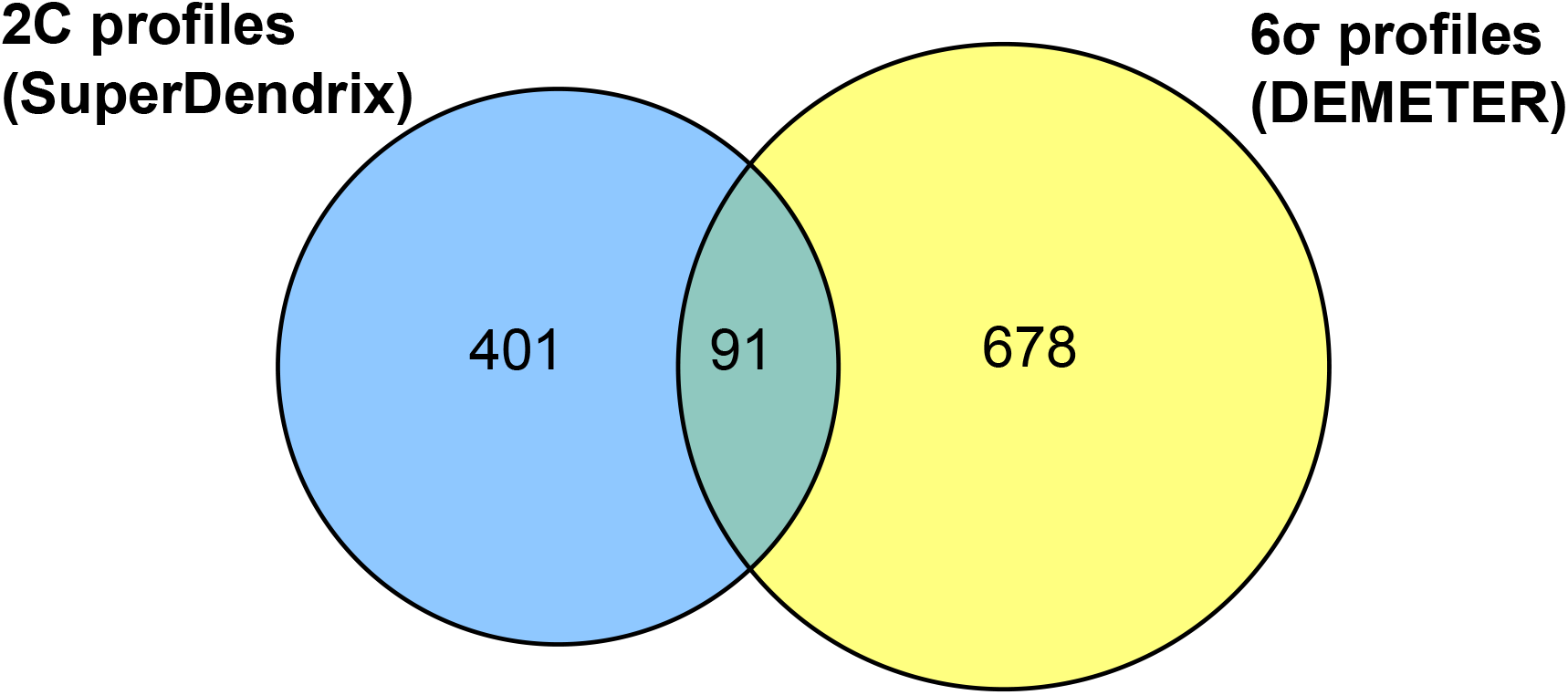
Comparison of differential dependencies found by SuperDendrix and DEMETER. Overlap between 492 differential dependencies identified by SuperDendrix from CRISPR dataset vs. 769 differential dependencies identified using 6*σ* cutoff in RNAi dataset from [15].

**Figure S13:**
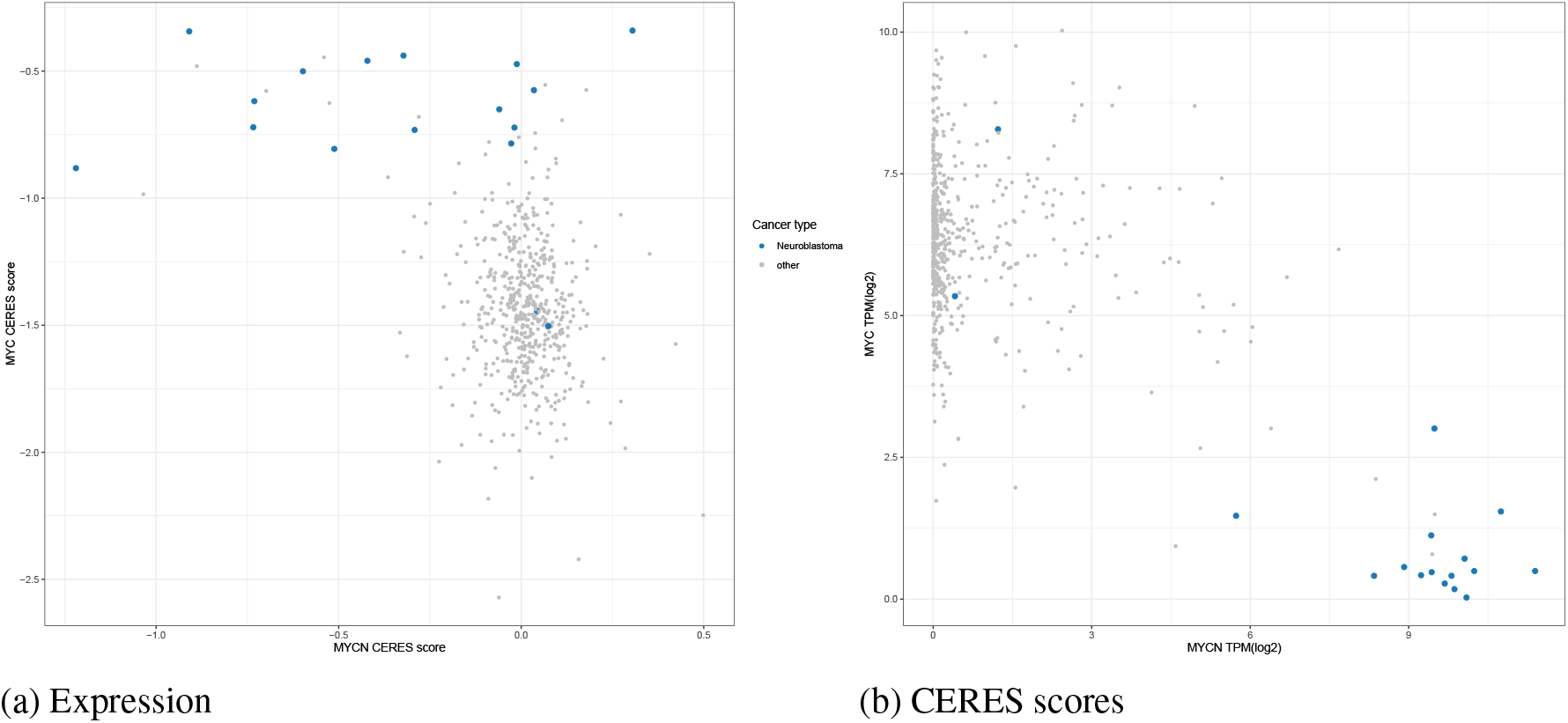
Dependencies on *MYC* and *MYCN* in neuroblastoma. (a) *MYC* is associated with decreased dependency in neuroblastoma. In contrast, *MYCN* is associated with increased dependency in neuroblastoma. (b) Consistent with these associations, *MYC* expression is high across cancer types except for neuroblastoma while *MYCN* expression is low except for the same cancer type.

## Supplementary Information

### Associations in Wnt pathway

SuperDendrix finds two associations in the Wnt pathway, including an association between `APC(I)` and `CTNNB1(A)` alterations and increased dependency in both *CTNNB1* and *TCF7L2* (Fig. S11). Alterations in `APC(I)` and `CTNNB1(A)` are mutually exclusive with no cell line containing both. Moreover, these dependencies are consistent with previously reported interactions in the Wnt pathway: when APC and correspondingly the *β*-catenin destruction complex are inactivated, CTNNB1 binds TCF7L2 in the nucleus and activates transcription of target genes [111].

### Comparison to other perturbation screen results

We compare the differential dependencies and alteration sets associated with these dependencies identified with our methods to the results of RNAi screening from Tsherniak et al. [15] and CRISPR-Cas9 screening from Behan et al. [16].

Tsherniak et al. [15] identified 6*σ* genes and associated genomic markers of these differential dependencies using RNAi screens of 501 cancer cell lines as part of the DepMap project. This analysis is distinct from ours in terms of the perturbation assay (RNAi instead of CRISPR) and score (DEMETER [15] instead of CERES), and in that Tsherniak et al. consider copy number aberrations and gene expression data – in addition to mutations – as potential genomic markers. There are 313 cell lines shared between the RNAi and CRISPR datasets.

We first compare in terms of differential dependencies. Tsherniak et al. [15] analyzed 6,305 profiles that pass quality control and identified 769 6*σ* genes. 91 of these 6*σ* profiles are also among the 492 2C differential dependencies (Fig. S12). Despite the small number of overlaps, the two sets of differential dependency profiles represent similar classes of proteins. In particular, both sets are significantly enriched for GO molecular functions such as DNA binding and protein kinase activity. They also contain similar proportion of CGC genes (Tsherniak et al. 6*σ*: 12.2% *P <* 0.01, 2C: 18.7% *P <* 0.01). Genes that are unique to each set also capture similar GO molecular functions including nucleotide binding, protein binding, and G protein-coupled receptor activity and are both significantly enriched for CGC genes (*P <* 0.01).

We next compare our results with Tsherniak et al. [15] in terms of biomarkers for differential dependency. Tsherniak et al. used a random forest-based approach to identify genomic features that are predictive of differential dependency, which they referred to as “marker dependency pairs” (MDPs). Using mutations, copy number aberrations, and gene expression, Tsherniak et al. [15] found MDPs for 426 of the 769 6*σ* profiles in the RNAi data. However, only 10 of these correspond to mutation driven biomarkers. In contrast, SuperDendrix found significantly associated mutation sets in 32 of 492 2C differential dependencies in the CRISPR data. 5 biomarker associations (mutation driven) are identified by both methods. These include well-known associations such as oncogene addictions of *BRAF*, *NRAS*, and *KRAS*. Interestingly, associations identified only by SuperDendrix include those with strong evidence, such as *RAF1* dependency on *KRAS* or *NRAS* mutations, *STAG1* dependency on *STAG2* mutations, and *NFE2L2* dependency on *KEAP1* mutations.

As part of DepMap project, Behan et al. [16] independently conducted genome-wide CRISPR-Cas9 loss-of-function screens in 324 cancer cell lines. From a total of 18,009 knockout genes, they identified 628 priority targets based on combination of gene knockout effect across cell lines and their associations to biomarkers (single nucleotide variants, copy number variations, and microsatellite instability status). 147 of the priority targets are also among the 492 2C profiles from SuperDendrix. The two sets of genes are significantly enriched for GO molecular functions such as DNA binding, protein binding, and transcription regulator activity. They also contain similar proportion of CGC genes (priority targets:15.9% *P <* 0.01, 2C: 18.7% *P <* 0.01).

Behan et al. analyzed associations of gene knockout effects with genomic biomarkers within each cancer type using ANOVA. Associations that occur across multiple cancer types were aggregated and re-tested using a *t*-test across all cell lines. We compare our results to their associations to SNVs considering all cell lines since we do not test for cancer-type-specific biomarker associations. Behan et al. identified a total of 41 significant biomarker associations (*FDR* ≤ 0.20) in 30 of the 628 priority target genes. However, only 10 associations for 8 genes are with SNV biomarkers. 3 of these (KRAS-KRASmut, PIK3CA-PIK3CAmut, GRB2-KRASmut) are also identified by SuperDendrix.

Overall, we are able to explain 6.5% (32) of 2C differential dependencies (492) with alterations using SuperDendrix, 9 of which contain more than one alteration. In contrast, Tsherniak et al. [15] and Behan et al. [16] can each explain only 1.3% 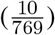 and 1.6% 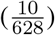 of their differential dependencies with mutation features. These findings indicate that our model, by searching for a set of approximately mutually exclusive alterations, has higher sensitivity for identifying associations between gene dependency biomarkers.

### Validation on the Sanger CRISPR-Cas9 screen data

We used the dataset from genome-wide CRISPR-Cas9 screens conducted as part of the Cancer Dependency Map at Wellcome Sanger Institute [16] to validate the associations identified by SuperDendrix from the Avana dataset of the Cancer Dependency Map at the Broad Institute.

First, we downloaded the Score dataset containing dependency scores computed from results of CRISPR screens across 324 cancer cell lines from the Project Score data portal^3^ and a list of mutations for the same cell lines from Cell Model Passports data portal^4^ [112]. We used quantile normalized log fold-changes as dependency scores and processed the mutation data using SuperDendrix OncoKB feature selection. We restricted our validation to 312 cell lines that contain at least one OncoKB mutation feature. For each association identified by SuperDendrix in the Avana dataset, we compared the dependency scores of cell lines containing at least one of the features with dependency scores of the cell lines without any feature. We found that 22 of the 32 associations between differential dependencies and mutations identified by SuperDendrix and 72 of the 117 associations between differential dependencies and mutations or cancer-type features identified by SuperDendrix are statistically significant in the Score dataset (*P* ≤ 0.05; Wilcoxon rank sum test). We find that many of the associations identified by SuperDendrix that did not validate in the Score dataset are in cancer types that were poorly represented in the Score dataset (Table S5).

### Decreased dependency on transcription factors

Three of the 41 transcription factors with cancer-type specific differential dependencies, *MYC*, *THAP1*, *TP53*, show *decreased dependency* in specific cancer types. For example, we find decreased dependency on *MYC* in neuroblastoma. Interestingly, neuroblastoma has increased dependency on *MYCN* – a well-known oncogene for neuroblastoma [113, 114] – another member of the same family of *MYC* genes (Fig. S13(a)). *MYCN* is highly expressed in a subset of tissue including forebrain and hindbrain whereas *MYC* expression is ubiquitous [114]. Previous studies have suggested that MYCN and MYC may be redundant, and that MYCN can compensate MYC activity [114, 115]. In fact, we observe that MYC expression is high across most cancer types in the dataset, with neuroblastoma a prominent exception; at the same time, MYCN expression is low across most cancer types except for neuroblastoma (Fig. S13(b)).Within neuroblastoma, expression of MYC and MYCN is approximately mutually exclusive, with most neuroblastoma cell lines expressing MYCN but not MYC. Based on these, we hypothesize that our result of the decreased dependency on *MYC* in neuroblastoma is because of its low expression in neuroblastoma, a result of redundant function and compensatory activity by MYCN. This result is consistent with the expected relationship between gene expression and dependency: cell lines in which a gene is not expressed will not be affected by knockout of that gene. As another example, we find decreased dependency on *THAP1* in leukemia and lymphoma. THAP1 is known as a pro-apoptotic factor involved in regulating endothelial cell proliferation and linking PAWR to promyelocytic leukemia (PML) nuclear bodies (NB) [116, 117]. Interaction of PAWR and PML has been reported to trigger apoptosis [118]. Furthermore, PML is a tumor suppressor primarily expressed in blood vessels and a negative regulator of cell survival pathways [116, 118]. These reports on lineage-specificity and function of THAP1 and PML suggest that knocking out *THAP1* which leads to loss of PML function resulted in decreased dependency or even prolonged cell survival in leukemia and lymphoma. We also find decreased dependency on *TP53* in rhapdoid, kidney cancer, and skin cancer. A possible explanation for decreased dependency on *TP53* is its wild-type function as a tumor suppressor. A previous study reports that knocking out *TP53* in cells with functional wild-type *TP53* where p53 acts as a tumor suppressor will induce growth advantage in those cells [119]. In our results, we noticed that many of the rhapdoid, kidney cancer, and skin cancer cell lines contain wild-type *TP53*. We thus tested for association between decreased dependency on *TP53* and `TP53(WT)` as an additional feature using SuperDendrix. In fact, SuperDendrix identified a significant association between them (*P* ≤ 0.001), confirming that this is a decreased dependency conferred by inhibiting tumor suppressor activity of p53 in *TP53* wild-type cell lines as suggested by [119].

### UNCOVER model

Although the underlying integer linear programming formulations differ, the optimization model of UNCOVER is similar to the SuperDendrix weight. Using the same notation from the SuperDendrix weight, the UNCOVER objective function is the following:

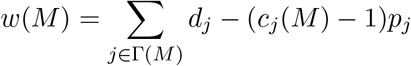

This function consists of two terms, *d_j_* and *p_j_*, that represent association to phenotype and penalty for co-occurring alterations. While UNCOVER uses the same linear term as SuperDendrix for biomarker-phenotype association, it uses a penalty term that has different values depending on the sign of the phenotype score. Specifically, if the phenotype score *d_j_* in cell line *j* is positive, then UNCOVER sets the penalty *p_j_* to the *average* of the positive phenotype scores; alternatively if the phenotype score *d_j_* in cell line *j* is negative then UNCOVER sets the penalty *p_j_* to be the absolute value of the score.

### Data preprocessing for UNCOVER

We use the same CERES data and alteration data^5^. For consistency, we process the phenotype scores and alteration features as described in UNCOVER. For phenotype scores, we normalize CERES scores in each knockout profile by subtracting the mean of the profile for each score and dividing the result by standard deviation of the profile. Since mixture model is a procedure specific to SuperDendrix pipeline, we run UNCOVER on 1,730 6*σ* profiles. For alterations, we combine nonsense, frameshift, and missense mutations of each gene as a single alteration feature. A gene alteration in a cell line exists if at least one of nonsense, frameshift, or missense mutation is found in that cell line. This results in 9,987 alteration features occurring across 554 cell lines.

## Supplementary Tables

All of the supplementary tables are attached in a supplementary information file.

Table S1: **Information on** 6*σ* **differential dependencies from CERES scores.**

Table S2: **Results of fitting t-mixture models to CERES dependency profiles.**

Table S3: **Enrichment for GO molecular functions in 2C profiles.**

**Table S4:**
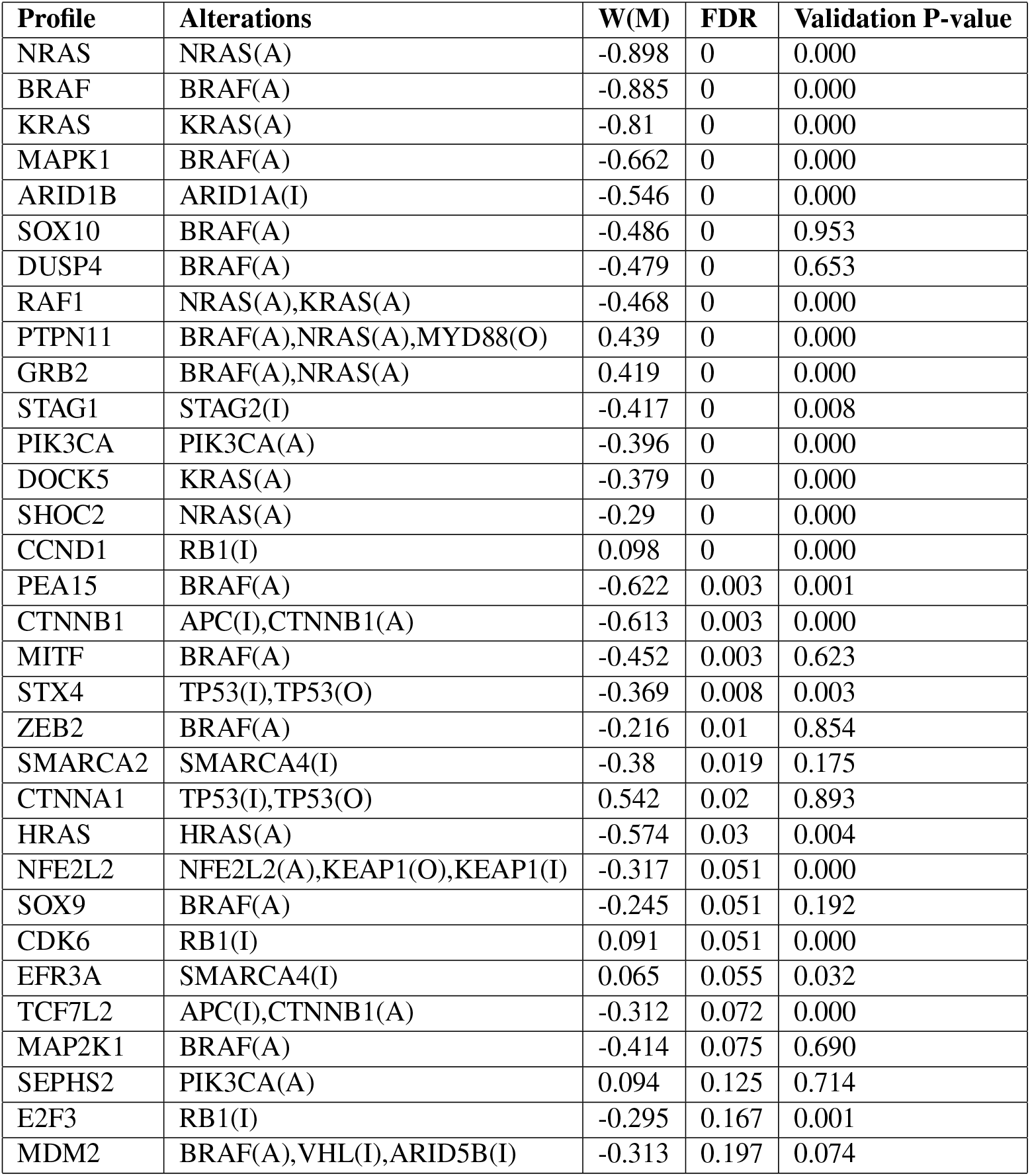
Associations between differential dependencies and sets of alterations identified by SuperDendrix. We rank results by FDR and the absolute value of SuperDendrix weight, 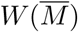.

Table S5: **Number of cell lines for each cancer type in the Avana and Score datasets.**

**Table S6:** Associations between differential dependencies and sets of alterations/cancer types identified by SuperDendrix. Same format as Table S4

| Profile | Alterations and cancer types | W(M) | FDR | Validation P-value |
| --- | --- | --- | --- | --- |
| NRAS | NRAS(A) | -0.898 | 0 | 0 |
| BRAF | BRAF(A) | -0.885 | 0 | 0 |
| MYOD1 | Rhabdomyosarcoma | -0.872 | 0 | NA |
| ISL1 | Neuroblastoma | -0.858 | 0 | 0 |
| KRAS | KRAS(A) | -0.81 | 0 | 0 |
| SOX10 | Skin | -0.781 | 0 | 0.103 |
| MEF2C | Multiple myeloma,Leukemia | -0.774 | 0 | 0.003 |
| MYB | Leukemia | -0.755 | 0 | 0.755 |
| ZEB2 | Leukemia,Skin,Ewings sarcoma,Rhabdomyosarcoma | -0.75 | 0 | 0.182 |
| MITF | Skin | -0.745 | 0 | 0.023 |
| GFI1 | Leukemia | -0.736 | 0 | 0.036 |
| IRF4 | Multiple myeloma,Lymphoma,Skin | -0.721 | 0 | 0.052 |
| POU2AF1 | Multiple myeloma,MEF2B(A) | -0.713 | 0 | 0.004 |
| HERPUD1 | Multiple myeloma | -0.699 | 0 | 0.006 |
| IKZF1 | Leukemia,Lymphoma,Multiple myeloma | -0.697 | 0 | 0 |
| HAND2 | Neuroblastoma | -0.689 | 0 | 0 |
| IGF2BP1 | Ewings sarcoma,Neuroblastoma | -0.682 | 0 | 0.002 |
| PRDM1 | Multiple myeloma | -0.681 | 0 | 0.251 |
| MAPK1 | BRAF(A) | -0.662 | 0 | 0 |
| FLI1 | Ewings sarcoma,Leukemia,Multiple myeloma | -0.661 | 0 | 0.249 |
| PAX8 | Kidney,Endometrial,Ovarian | -0.661 | 0 | 0 |
| ATP1B3 | Lymphoma,Multiple myeloma,Leukemia | -0.633 | 0 | 0 |
| PEA15 | BRAF(A) | -0.622 | 0 | 0.011 |
| SPDEF | Breast | -0.587 | 0 | 0.962 |
| TP63 | Head and Neck,Bladder,Esophageal | -0.524 | 0 | 0 |
| CCND3 | Leukemia,Lymphoma | -0.486 | 0 | 0.85 |
| TCF7L2 | Colon,Gastric | -0.477 | 0 | 0 |
| DUSP4 | Skin | -0.469 | 0 | 0.003 |
| RAF1 | NRAS(A),KRAS(A) | -0.468 | 0 | 0 |
| MYC | Neuroblastoma | 0.468 | 0 | 0 |
| SPI1 | Leukemia | -0.461 | 0 | 0.393 |
| PIK3CA | PIK3CA(A),Multiple myeloma | -0.453 | 0 | 0 |
| PTPN11 | BRAF(A),NRAS(A),MYD88(O) | 0.439 | 0 | 0 |
| GRB2 | BRAF(A),NRAS(A) | 0.419 | 0 | 0 |
| NAMPT | Leukemia,Multiple myeloma,Breast,Lymphoma | -0.413 | 0 | 0.04 |
| FERMT2 | Skin,Brain,Kidney,Ovarian,Sarcoma | -0.411 | 0 | 0 |
| LDB1 | Leukemia,Neuroblastoma | -0.36 | 0 | 0 |
| CDK6 | RB1(I),Breast,Endometrial,Ewings sarcoma,Ovarian | 0.351 | 0 | 0 |
| SEPHS2 | Head and Neck,Colon,Brain,Esophageal,Gastric | 0.346 | 0 | 0 |
| UMPS | Leukemia,Lymphoma,Multiple myeloma,Breast | -0.341 | 0 | 0.019 |
| SHOC2 | NRAS(A),Brain | -0.339 | 0 | 0 |
| GPX4 | Colon,Head and Neck,Esophageal,Skin,Gastric | 0.324 | 0 | 0 |
| IRS2 | Ewings sarcoma,Neuroblastoma | -0.321 | 0 | 0.004 |
| CFLAR | Skin,Brain,Neuroblastoma,KIT(O),Ewings sarcoma | 0.316 | 0 | 0.001 |
| PSMB5 | Leukemia,Lymphoma,SMARCB1(I) | 0.296 | 0 | 0.124 |
| CCND1 | RB1(I),Leukemia,Lymphoma,Rhabdomyosarcoma | 0.294 | 0 | 0 |
| SNAP23 | Brain,Neuroblastoma,Skin,Ewings sarcoma | 0.292 | 0 | 0 |
| EFR3A | SMARCA4(I),Neuroblastoma,Skin,Breast | 0.22 | 0 | 0 |
| GFPT1 | Leukemia,VHL(I) | -0.19 | 0 | 0.013 |
| DNM2 | Neuroblastoma | 0.187 | 0 | 0.015 |
| PGM3 | Brain,Lung NSCLC | 0.175 | 0 | 0 |
| TCF3 | Multiple myeloma,ID3(I),MEF2B(A) | -0.632 | 0.001 | 0.27 |
| RELB | Multiple myeloma | -0.588 | 0.001 | 0.005 |
| ARID1B | ARID1A(I) | -0.546 | 0.001 | 0.003 |
| SLC5A3 | Leukemia | -0.449 | 0.001 | 0.089 |
| STAG1 | STAG2(I) | -0.417 | 0.001 | 0.016 |
| DOCK5 | KRAS(A) | -0.379 | 0.001 | 0 |
| STX4 | TP53(I),TP53(O) | -0.369 | 0.001 | 0.177 |
| CCND2 | Multiple myeloma | -0.325 | 0.001 | 0.003 |
| TRPS1 | Breast | -0.51 | 0.002 | 0.003 |
| GRHL2 | Head and Neck,Bladder,Breast,Pancreatic | -0.377 | 0.002 | 0.502 |
| PTPN2 | Lymphoma | -0.588 | 0.002 | NA |
| DCAF5 | Rhabdoid | -0.55 | 0.002 | NA |
| BCL2 | Leukemia,Multiple myeloma,MEF2B(A),Neuroblastoma | -0.534 | 0.003 | 0 |
| NFATC2 | Skin | -0.43 | 0.003 | 0.668 |
| SMARCA2 | SMARCA4(I) | -0.38 | 0.003 | 0.189 |
| ATP1A1 | Neuroblastoma | 0.147 | 0.003 | 0.002 |
| ATP2C1 | Multiple myeloma | -0.349 | 0.004 | 0.005 |
| CTNNA1 | TP53(I),TP53(O) | 0.542 | 0.005 | 0.982 |
| TP53 | Rhabdoid,Kidney,Skin | 0.468 | 0.005 | 0.288 |
| LMO2 | Leukemia | -0.323 | 0.005 | 0.198 |
| PSTK | Colon,Esophageal,FLCN(I) | 0.177 | 0.005 | 0 |
| YRDC | Brain | 0.079 | 0.005 | 0 |
| SMAD7 | Multiple myeloma | -0.484 | 0.005 | 0.003 |
| RPP25L | Brain,Lymphoma,Rhabdomyosarcoma | -0.248 | 0.006 | 0 |
| TFAP2B | Neuroblastoma,Rhabdomyosarcoma | -0.551 | 0.006 | 0.006 |
| RAB6A | SMARCA4(I),Skin | 0.19 | 0.006 | 0.005 |
| MDM2 | Skin,Kidney,ARID5B(I) | -0.352 | 0.007 | 0.215 |
| MET | Brain | -0.29 | 0.007 | 0.005 |
| MARCH5 | Multiple myeloma,Leukemia | -0.288 | 0.007 | NA |
| GET4 | Kidney | -0.159 | 0.007 | 0.122 |
| BATF3 | Lymphoma,Multiple myeloma | -0.537 | 0.008 | 0.133 |
| MYCN | Neuroblastoma,Rhabdomyosarcoma | -0.466 | 0.009 | 0.013 |
| HRAS | HRAS(A) | -0.574 | 0.01 | 0.001 |
| FOXA1 | Breast | -0.394 | 0.01 | 0.612 |
| CHUK | Multiple myeloma,Lymphoma | -0.545 | 0.012 | 0.01 |
| LIN28B | Neuroblastoma,Ewings sarcoma | -0.444 | 0.012 | 0.019 |
| VCL | Brain | -0.237 | 0.012 | 0 |
| SOX9 | Skin,Colon,Gastric,Pancreatic | -0.479 | 0.012 | 0.056 |
| PAX3 | Rhabdomyosarcoma | -0.351 | 0.012 | NA |
| IGF1R | Multiple myeloma,Ewings sarcoma,Rhabdomyosarcoma | -0.322 | 0.012 | 0.58 |
| ARHGEF7 | KRAS(A),Head and Neck,Bladder,Liver | -0.415 | 0.016 | 0 |
| NFE2L2 | NFE2L2(A),KEAP1(O),KEAP1(I) | -0.317 | 0.016 | 0.005 |
| DHODH | Leukemia | -0.247 | 0.018 | 0.003 |
| THAP1 | Leukemia,Lymphoma | 0.219 | 0.019 | 0.205 |
| IKBKB | Multiple myeloma,Lymphoma | -0.607 | 0.02 | 0.006 |
| IRS1 | Multiple myeloma,Rhabdomyosarcoma | -0.414 | 0.021 | 0.104 |
| MAP2K1 | BRAF(A) | -0.414 | 0.021 | 0 |
| TFAP2C | Breast | -0.341 | 0.022 | 0.103 |
| ALG12 | Multiple myeloma,Lung | -0.49 | 0.031 | 0.015 |
| CTNNB1 | Colon,Gastric,Liver,Pancreatic | -0.646 | 0.031 | 0 |
| CAD | Leukemia,Multiple myeloma,Breast,Lymphoma | -0.304 | 0.031 | 0.009 |
| EGFR | Head and Neck | -0.226 | 0.032 | 0 |
| KCTD10 | TP53(I),TP53(O),Neuroblastoma,Rhabdoid | 0.25 | 0.032 | 0.44 |
| GCLC | Leukemia,Lymphoma,Multiple myeloma,GATA3(I) | -0.634 | 0.037 | 0 |
| AHCYL1 | Kidney,Liver | -0.172 | 0.037 | 0.251 |
| E2F3 | RB1(I) | -0.295 | 0.042 | 0 |
| JUN | Brain | -0.244 | 0.045 | 0 |
| SEC23IP | Kidney | -0.25 | 0.049 | 0.926 |
| HSPA13 | Head and Neck,Esophageal | -0.202 | 0.052 | 0 |
| PTK2 | Skin,Brain | -0.186 | 0.055 | 0 |
| PPAT | Leukemia | -0.151 | 0.065 | 0.008 |
| CHEK2 | Kidney | 0.29 | 0.075 | 0.497 |
| DMXL1 | Multiple myeloma,MEF2B(A) | -0.511 | 0.08 | 0.113 |
| MVK | Brain | 0.081 | 0.08 | 0.008 |
| FURIN | Multiple myeloma,Ewings sarcoma,Neuroblastoma,Leukemia,Pancreatic | -0.525 | 0.086 | 0.001 |
| TFAP2A | Skin | -0.317 | 0.086 | 0.92 |
| IPPK | Leukemia,Multiple myeloma,Lymphoma | -0.346 | 0.088 | 0.044 |
| PPCDC | Leukemia,Lymphoma,Multiple myeloma | -0.328 | 0.088 | 0.4 |
| SCAP | Neuroblastoma,ARID1B(I),Bone | 0.198 | 0.089 | 0 |
| FPGS | Leukemia,Breast,Lymphoma | -0.276 | 0.102 | 0.08 |
| PPARG | Bladder,Pancreatic | -0.442 | 0.106 | 0.399 |
| WDR4 | Head and Neck | -0.176 | 0.106 | 0.025 |
| FOSL1 | NF2(I),Head and Neck | -0.175 | 0.117 | 0 |
| HNF1B | Kidney,Endometrial,Liver | -0.405 | 0.119 | 0.071 |
| PARD6B | Kidney,Ovarian,Liver | -0.307 | 0.122 | 0 |
| SEPSECS | Colon | 0.106 | 0.125 | 0.109 |
| SMARCE1 | Rhabdoid | 0.122 | 0.136 | NA |
| SMARCB1 | Rhabdoid,Sarcoma,Kidney | 0.302 | 0.141 | 0.172 |
| EGLN1 | Skin,Ovarian | -0.337 | 0.15 | 0.321 |
| RFK | Leukemia,Lymphoma,PBRM1(O),EPHA7(O) | -0.329 | 0.158 | 0.001 |
| SLC4A7 | Lymphoma | -0.106 | 0.158 | NA |
| PPP1R12A | Leukemia,Neuroblastoma | 0.226 | 0.161 | 0.002 |
| ERBB2 | Head and Neck | -0.183 | 0.188 | 0.005 |
| ZBTB38 | Multiple myeloma | -0.343 | 0.192 | 0.004 |

Table S7: **Enrichment for GO molecular functions in significant cancer-type dependencies.**

Table S8: **Enrichment for GO molecular functions in non-transcription factors with significant cancer-type dependencies.**

Table S9: **List of 558 DepMap cell lines (2019Q1).**

Table S10: **Alteration dependencies from UNCOVER.**

Table S11: **Alteration or cancer-type dependencies from UNCOVER.**

## Footnotes

1 https://depmap.org/portal/download/

2 https://ndownloader.figshare.com/files/14941493

3 https://score.depmap.sanger.ac.uk/downloads

4 https://cellmodelpassports.sanger.ac.uk/downloads

5 https://depmap.org/portal/download/api/download/external?file_name=ccle%2Fdepmap-mutation-calls-9a1a.9%2Fdepmap_19Q1_mutation_calls_v2.csv

